# Flossing DNA in a Dual Nanopore Device

**DOI:** 10.1101/778217

**Authors:** Xu Liu, Philip Zimny, Yuning Zhang, Ankit Rana, Roland Nagel, Walter Reisner, William B. Dunbar

## Abstract

Solid-state nanopores are a single-molecule technique that can provide access to biomolecular information that is otherwise masked by ensemble averaging. A promising application uses pores and barcoding chemistries to map molecular motifs along single DNA molecules. Despite recent research breakthroughs, however, it remains challenging to overcome molecular noise to fully exploit single molecule data. Here we present an active control technique termed “flossing” that uses a dual nanopore device to trap a protein-tagged DNA molecule and perform up to 100’s of back-and-forth electrical scans of the molecule in a few seconds. The protein motifs bound to 48 kb *λ*DNA are used as detectable features for active triggering of the bidirectional control. Molecular noise is suppressed by averaging the multi-scan data to produce averaged inter-tag distance estimates that are comparable to their known values. Since nanopore feature-mapping applications require DNA linearization when passing through the pore, a key advantage of flossing is that trans-pore linearization is increased to >98% by the second scan, compared to 35% for single nanopore passage of the same set of molecules. In concert with barcoding methods, the dual-pore flossing technique could enable genome mapping and structural variation applications, or mapping loci of epigenetic relevance.

## Introduction

A solid-state nanopore refers to a nanoscale hole formed in a solid-state membrane^1^ or at the tip of a glass pipette.^2^ A voltage-clamp amplifier supplies a voltage that is concentrated across the pore to electrically measure the trans-pore ionic current. When the voltage captures a single DNA and drives it through the pore, the passing DNA produces a transient blockade in the current that contains information about the molecule’s chemical, conformational and topological state. Nanopore sensing offers a simple and high-throughput electrical read-out ^3^ with an instrument that can have a small foot-print at low cost. ^4^

Recent research has showcased the potential of nanopores to detect molecular features along a DNA carrier strand, including proteins such as anti-DNA antibodies^5^ and streptavadin,^6,7^ single-stranded versus double stranded regions of a molecule, ^8^ DNA-hairpins^9,10^ and apatamers.^11,12^ Potential applications range from digital information storage, ^9,10^ multiplexed sensing, ^9,11^ and genomic and/or functional genomic applications including genome mapping^13^ and epigenetics. ^14,15^ Solid-state pores in particular can target a more diverse analyte pool than protein pores^16^ (e.g. dsDNA, proteins, protein-DNA complexes, nucleosomes^17^) and thereby give access to a broad range of single-molecule applications.

A key challenge in performing multi-locus sensing of motifs along DNA is the inherent sensitivity of single-molecule systems to noise. Unwanted conformations/topologies and molecular fluctuations (both equilibrium and non-equilibrium in nature) create systematic and random distortions in the electrical signal pattern of motifs resolved by the sensor. For example, closely spaced features along DNA cannot always be resolved in a given single molecule read even with state-of-the-art measurements performed with 5 nm diameter nanopores,^10^ requiring multiple independent reads from identical copies of different molecules to confidently resolve the features. In addition, the stochastic nature of the translocation process gives rise to broad distributions^18,19^ in tag spacings measured across a molecular ensemble;^7^ these broad distributions necessitate averaging over additional molecules to obtain precise spacing estimates. A related challenge is providing independent genomic distance calibration along individual single-molecule reads, so that sensor output can be linked to sequence position without a priori knowledge of the distance between motifs. While optics can provide high-resolution spatiotemporal data,^20,21^ nanopores can only infer spatial information implicitly from temporal data. In order for nanopore technology to achieve its full potential, it is essential that single-molecule reads have sufficient quality (e.g., contain sufficiently low systematic and random errors), so that the requirement for further ensemble-level averaging over different molecules is minimized or eliminated. The technology can then be applied to the complex, heterogeneous samples reflective of applications where every molecule may have a different number of bound motifs possessing a distinct spatial distribution. Nanochannel technology, for example, harnesses the physics of nanofluidic polymer confinement to both linearize DNA and suppress molecular fluctuations in barcoded molecules, giving rise to powerful genome-scale applications in the area of DNA optical mapping.^22,23^

Here we show that highly accurate spatial information that is correlative with motif binding can be obtained from a *single* labeled carrier dsDNA strand via repeated back-and-forth scanning of the molecule trapped in a nanopore device. Using a new active control technique termed “flossing,” we are able to perform up to 100’s of scans of a given trapped molecule within a few seconds. Flossing is showcased here using a model system consisting of a 48.5 kbp double-stranded *λ*-DNA with a set of chemically incorporated sequence-specific protein tags. Our approach complements existing carrier-strand DNA nanopore technology by enhancing the quality of information that can be extracted from a single trapped molecule. By taking a large number of statistically equivalent scans, we reduce stochastic fluctuations through scan averaging, and show mean tag spacing estimates that are comparable to known inter-tag distances, even while tolerating missed tag(s) within a subset of scans.

While prior work leveraged control logic for automated re-capture and sensing of molecules with single nanopores,^24,25^ our approach is distinctive in a few ways. First, our device employs a dual-pore architecture.^26–28^ By having the dual pores sufficiently close, our prior work shows the DNA can be captured simultaneously by both pores and exist in a ‘tug-of-war’ state where competing electrophoretic voltage forces are applied at the pores. ^28^ During tug-of-war, the molecule’s orientation and identity are maintained, the molecule speed is regulated to facilitate tag sensing, and the likelihood of the molecule finding a linearized conformation through the pore is increased to 70% compared to 30% with single pore data. An advantage of flossing is that linearization through the pore is further increased to >98% by the second scan, which in turn increases the throughput of nanopore feature-mapping applications.

Another distinct feature of the presented approach is that the active controller cyclically modulates the voltage at one pore by a real-time feedback on the sensing current of the other pore. Specifically, during control, the cyclical application of unbalanced competing voltage forces are used to drive the molecule’s motion in one direction and then, after real-time detection of a set number of tags, in the reverse direction, thereby embodying the concept of DNA “flossing.” Previously, the original concept of flossing DNA in a nanopore was envisioned with single protein pores, and only by creating stoppers at the end of the DNA to prevent escape.^29^ This approach facilitated interesting research^30,31^ but did not provide a means of speed control or of mineable data generation *during* the molecule’s motion in either direction. Coupling the DNA to a stage has produced both speed control and mineable data generation during the molecule’s motion through a solid-state pore, but at the price of complex instrumentation, higher sensing noise and lower throughput.^32^ In the presented flossing control method, an interrogated DNA molecule can be ejected from the pores and a new DNA captured with the same throughput and ease of any single-nanopore based assay, with no tethering of the molecule required.

## Results and Discussion

### The DNA Flossing Concept

Figure 1 introduces the general flossing concept, showing pictorially and with actual recorded data the cyclical bidirectional scans of a co-captured molecule in a dual nanopore device. Our dual nanopore device was fabricated as previously described,^33^ with voltages V_1_ and V_2_ that can be independently applied at pore 1 and 2, respectively. Two currents (I_1_ and I_2_) can also be independently measured at the pores. The tagged reagent features monovalent streptavidin (MS) proteins bound along the DNA.

**Figure 1:**
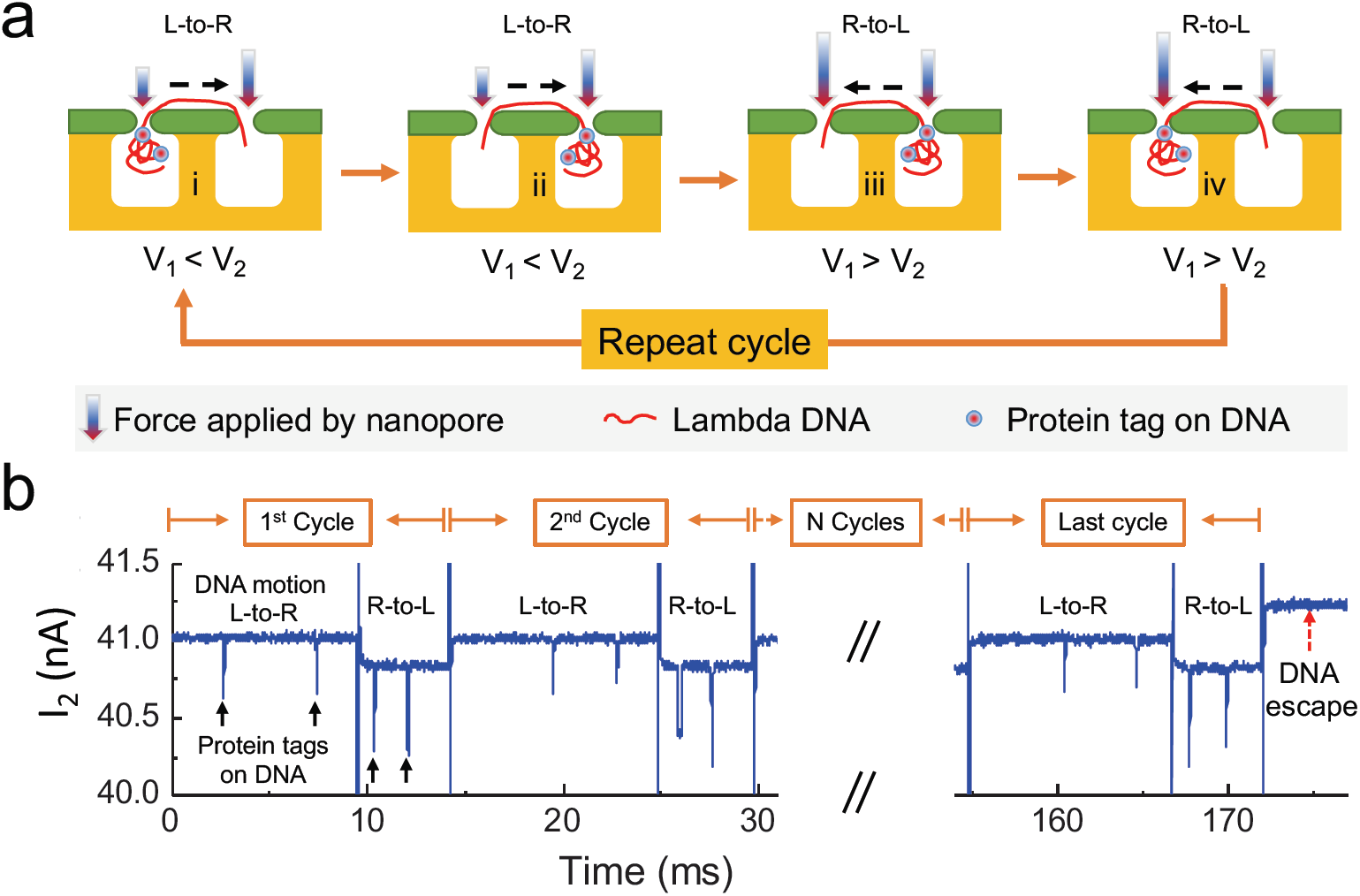
Flossing DNA with competing voltage forces in a dual pore device. **a** After DNA co-capture, the DNA molecule will be threaded from left-to-right (L-to-R) using a voltage V_1_ <V_2_, with V_1_ and V_2_ the voltages across pore 1 (left) and pore 2 (right), respectively. A single transit of DNA motion during this fixed polarity period is called a “scan.” After automated detection of a predefined number of tags, the direction of DNA motion is reversed with a voltage V_1_ >V_2_ triggered to move the molecule from right-to-left (R-to-L), giving rise to a second scan. The process is repeated in cyclical fashion until the molecule randomly exits the co-capture state. **b** A recorded multi-scan current trace I_2_ from pore 2, using logic for which the predefined tag detection number is 2, after which the controller triggers the change in direction. The signal from 30-150 ms is truncated for visualization.

Figure 1a illustrates each step in a multi-scan cycle. Since the motion control is bidirectional, we define a single transit with fixed direction as a “scan” and two sequential scans of reversed polarity as a “cycle.” By convention, we define the left pore as pore 1 and the right pore as pore 2. During the multi-scan control logic sequence, V_2_ across pore 2 is kept constant while V_1_ is modulated in step-wise fashion. The signal I_2_ is monitored in real-time for tag-related events as a logic trigger for V_1_ changes, as described next.

The flossing control logic begins by initially co-capturing a tagged DNA in both pores to reach the tug-of-war state. Once co-capture occurs, a lower voltage is applied across pore 1 than pore 2 (V_1_ < V_2_) to direct the molecule motion towards pore 2 (left-to-right, or “L-to-R”). While monitoring I_2_, the control logic readjusts the voltage at pore 1 so that V_1_ > V_2_ after a set number *N* of tags were detected translocating through pore 2. The readjustment directs the molecule motion back towards pore 1 (R-to-L). After detecting the same set number *N* of tags translocating through pore 2, the logic resets V_1_ < V_2_ to initiate another L-to-R scan, and thus a new cycle begins. Figure 1b shows a recorded example of the first two cycles and the final cycle of the I_2_ signal for which the tag number setting was *N* = 2. In this example, the multi-scan cycle begins with the molecule moving L-to-R for the first 9 ms and continues until 172 ms when the DNA escapes just after a V_1_ modulation (escape modes are discussed below and in detail in the SI). Details on the set of dual-pore chips and voltage settings used in this paper are provided in Tables S1-S2. Having described the general flossing concept, we next present the method in greater detail and results obtained using the method.

### Initializing Tug-of-War and Identifying Scanning Voltages

The control logic that automates the flossing process shown in Figure 1 is described here in detail. The flossing method first uses active control to automate initializing co-capture and tug-of-war on a single molecule. The control logic was run in real-time (MHz clock rate) on a Field Programmable Gate Array (FPGA). We modified our previously designed tug-of-war control process^28^ to permit loading reagent in the common fluidic chamber above the two nanopores and to screen out short-fragments (SI Section 2, Fig. S1). Once co-capture is achieved the competing voltage forces at V_1_ and V_2_ lead to a tug-of-war and reduce the DNA speed during sensing. The mean durations of all co-captured events are computed at each of a set of different V_1_ values while keeping V_2_ constant, showing a bell curve with the peak revealing the force-balancing voltages that maximize co-capture duration. Figure S1c shows an example device with peak mean duration of 110 ms with V_1_ and V_2_ both set to 500 mV. This data is generated with bare DNA that has no tags.

While capturing a molecule in dual pore tug-of-war introduces speed and conformational control, bare double-stranded DNA offers no detectable features with which to monitor the molecule’s motion. To enable *in-situ* feature monitoring, we developed MS-tagged DNA as a model reagent, with up to 7 sites tagged on the DNA (SI Section 3, Fig. S2). The tag features are used to make inferences on the speed and direction of each scan, which enables identifying the scanning voltages as described next.

The scanning voltages are found after the force-balancing voltages that maximize co-capture duration have been identified for a given chip. Specifically, the scanning voltage values for V_1_ are chosen above and below it’s force balancing value in order to promote unperturbed DNA motion toward pore 1 and pore 2, respectively, while keeping V_2_ at it’s force balancing value. The V_1_ scanning values are chosen heuristically, using the following guidance. A working scan speed should enable robust sensing of tag blockades at the recording bandwidth (i.e., not too fast), while ensuring sufficiently high Peclet numbers so that translocation time distributions are well-defined (i.e., not too slow). That is, broad translocation time distributions increase the probability of fluctuations, which can undermine the inter-tag time-to-distance mapping objectives described in a later section. In practice, achieving a not-too-fast and not-too-slow scan speed is achievable for a broad range of scanning V_1_ voltages. For the data in this paper, that range is as low as 150 and as high as 500 mV away from the force-balancing voltage (Table S2). Figure S2 shows an example tug-of-war experiment with MS-tagged DNA at a working scan speed, for which V_1_ = 200 mV and V_2_ = 500 mV, with a typical current traces for I_1_ and I_2_ from one representative event (Fig. S2c). There are three downward spikes in both I_1_ and I_2_, indicating three resolvable tags along the DNA. The V_1_ = 200 mV value that promotes L-to-R motion is 300 mV below the force-balancing value (500 mV).

### Representative Flossing Event with Protein-Tagged DNA

Once the scanning voltage values are identified, the full flossing multi-scan logic can be applied. The full details of the FPGA-implemented logic are provided in supplementary materials (SI Section 4, Figure S3). The output of the logic is conveyed here by representative flossing data (Figure 2). Figure 2a shows the full signal trace of I_1_, V_1_, I_2_ and V_2_ from a typical flossing event. We kept V_2_ = 300 mV unchanged within the event. We set V_1_ = 100 mV for L-to-R movement and V_1_ = 850 mV for R-to-L movement. Note that the I_2_ baseline varies when V_1_ is adjusted even though V_2_ is constant. This effect is caused by cross talk between pore 1 and pore 2 that was previously characterized in Liu *et al.*^28^ The detectability of tags relative to the DNA baseline in I_2_ is still robust, despite the baseline change. As detailed in the previous section, we chose sufficiently large voltage differentials between V_1_ and V_2_ to promote controlled motion in each direction during sensing.

**Figure 2:**
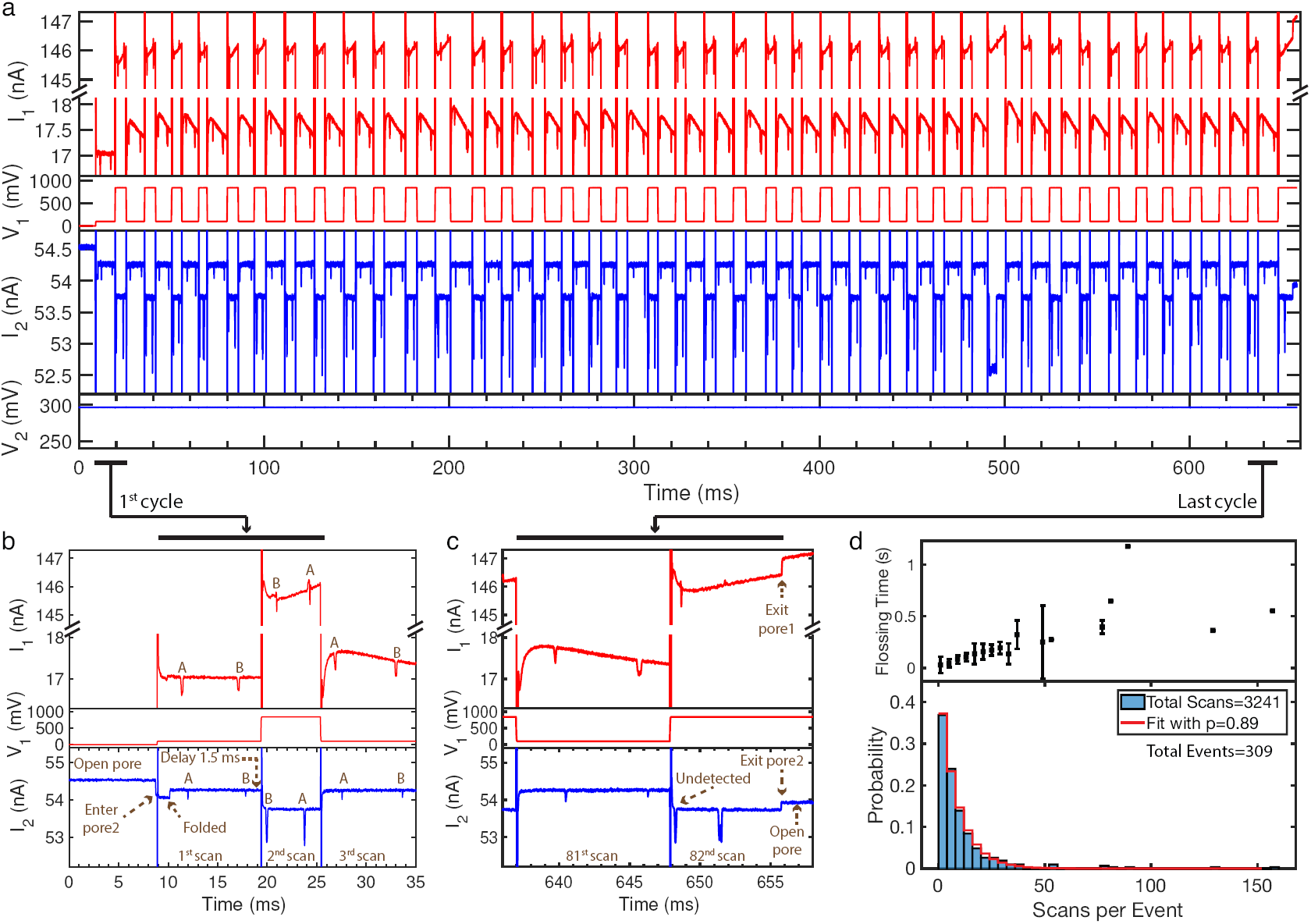
Representative dual current signals and scan count statistics generated during a flossing experiment with MS-tagged DNA. **a** Full signal traces for I_1_, V_1_, I_2_ and V_2_ are shown for a representative multi-scan flossing event. The vertical-axis break in the I_1_ signal permits vertical-scale zooming on the low and high ranges during the lower and higher V_1_ values. **b** Zoom in of the 1st cycle where two-tag logic shows resolvable tags A and B in both signals. **c** Zoom-in of the 41st and last cycle, showing the end of co-capture due to an undetected tag. **d** The total flossing time (mean ± standard deviation) and probability distribution versus scan counts across all co-captured events for the device used (bin width = 4). The red line on the probability data is the fitted model equation (1), with *p* = 0.89 the probability of correctly detecting two tags in each scan. The chip used had a pore-to-pore distance of 0.61 *µ*m, 27 nm pore 1 diameter, and 25 nm pore 2 diameter (Chip C, Table S1).

Figure 2b,c give a magnified view of the signal for the first cycle that comprises the 1st and 2nd scans, and the last cycle comprising the 81st and 82nd scans. By convention, since the multi-scan logic starts in the L-to-R direction, the odd scans correspond to L-to-R movement and the even scans R-to-L movement. The event in Figure 2a includes 41 cycles and 82 scans total, all in less than 0.66 sec. As depicted in Figure 1, the FPGA logic was designed to switch V_1_ once two tags were detected within I_2_. The FPGA detects a tag when I_2_ falls at least 70 pA below the untagged DNA baseline for at least 0.012 ms (based on the distribution in Figure S2f). The data in Figure 2 shows two tags, A and B, in both I_1_ and I_2_ when the molecule moved L-to-R from 9 to 19 ms. Following a 1.5 ms-delay after tag B was detected, the FPGA set V_1_ = 850 mV, driving the molecule to move R-to-L from 19 to 25 ms. The same tags, B and A, were detected in both I_1_ and I_2_ in reverse order. The same logic continued until the FPGA failed to detect tag B in the last cycle (Figure 2c), which was caused by the tag appearing too close to the voltage change for the FPGA to detect it.

The total flossing time and distribution of the number of scans per event are shown in Figure 2d for a total of 309 flossing events in an experiment, including the 82 scan event in Figure 2a. Total flossing time increases with more scans, while all events terminated in less than a few seconds, even for the largest scan count of 157 for this data. While individual scans last 5-10 ms (Figures 1b and 2b-c), the total time data shows a significant increase (up to 100X more) in time spent interrogating each molecule by using the flossing method.

From the probability plot in Figure 2d, 37% of the events had less than 5 scans, and events with higher scan count are less likely. We examined the probability *P* (*n*) of seeing a specific total number of scans *n*, where the probability of any intermediate scan has correct detection probability *p* and missed detection probability (1 −*p*). An event with *n* total scans indicates the system successfully catches the initial (*n* − 1) scans but fails to catch the *n*th scan. Thus its probability *P* (*n*) is

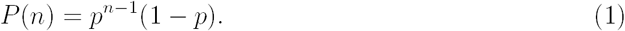

Fitting the data to equation (1) results in *p* = 0.89 for this specific data set (Figure 2d). To determine why the molecule exits co-capture, we studied the last cycles and found four common cases: missing a tag in the *n*th scan (Figure S4a); a false positive spike detected in the (*n* − 1)st scan (Figure S4b); a false negative spike in the *n*th scan (Figure S4c); and the molecule exits the pore during the FPGA delay state (Figure S4d). A discussion on how to increase the total number of scans per event can be found in SI section 5. Naturally, changing the voltage settings will affect event duration between tags, which in turn will affect tag detection probabilities.

A dependence of tag amplitude on scan direction is observed in Figure 2a-c, with MS tags showing relatively shallower and faster spikes when passing through the pore with the higher voltage of the two. Seeing a faster and shallower tag event at higher voltage is consistent with single pore results, and is in part an artifact of the low-pass filter (10 kHz bandwidth) preventing the tag events from hitting full depth (i.e., the faster the event, the shallower).

A multi-scan experiment using a three tag triggering setting was also performed (Figure S5). It is harder to get a higher scan count using three tag triggering than with two. In part, this arises because fewer molecules (<20%) show three or more tags (Figure S2e), which is a limitation of our current reagent preparation methods. Also, even when the system co-captures one molecule with three tags in both pores, the probability of correct detection of all three tags is lower (Figure S5c), and failing to detect any one of the tags moves the molecule to a new region, thereby lowering the scan count for the originally scanned three tag region. To generate more data with higher scan counts, we therefore focused the experiments on two-tag triggering in this initial presentation.

### Flossing Increases the Fraction of DNA with Mappable Data

Nanopore feature-mapping applications require DNA linearization when passing through the sensing pore. As such, we explored the fraction of DNA that can be linearized by the flossing technique, where linearization refers to the removal of DNA folds that are initially in the pore when co-capture is initialized. By example, the molecule in Figure 2a-c was partially folded at around 10 ms in the 1st cycle (Figure 2b), which was eliminated in the 2nd cycle and thereafter, demonstrating the tendency of tug-of-war control to induce and maintain DNA linearization. We examined the probability of complete linearization over the scanning cycle through pore 2 as a function of scan number to see if the trend in Figure 2a-c was representative of the population. Indeed, Figure 3 shows that the probability of linearization is increased to 98% by the second scan. In the data, a folded event is identified if the current blockage is larger than 1.5 times the unfolded blockage amount, and lasts more than 180 *µ*s. Figure 3a,b show representative single pore and multi-scan events with observable folding examples.

**Figure 3:**
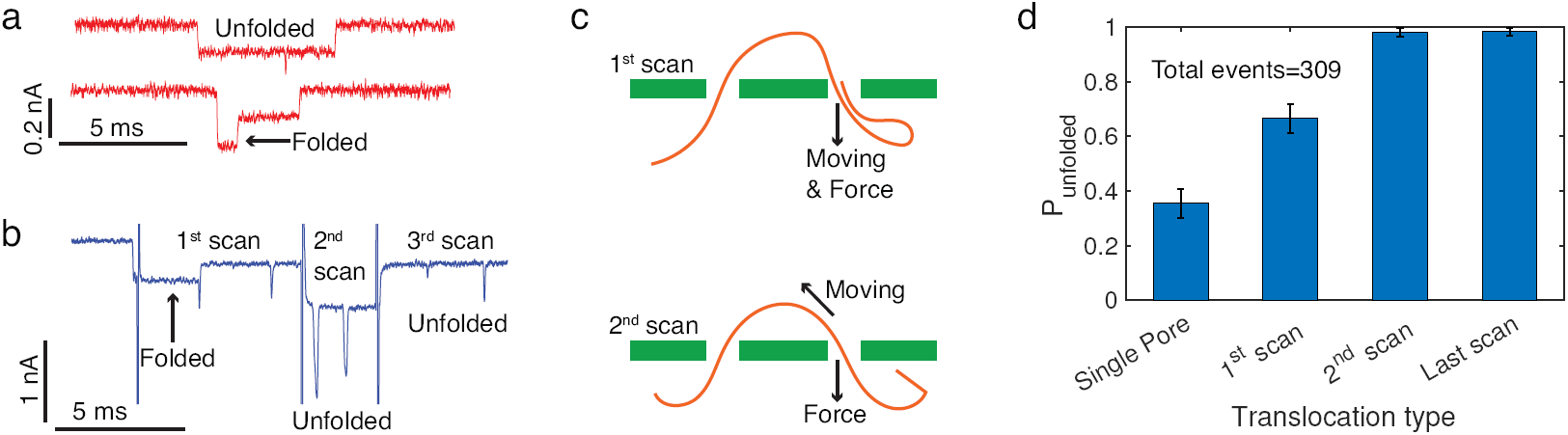
Flossing increases linearization of DNA in dual pore device. **a** Typical I_1_ traces of single pore events, including both unfolded folded examples. Only single pore events that resulted in eventual co-capture were included in subsequent probability calculations (pre-i step events in Figure S1). **b** Typical I_2_ trace of a multi-scan event in which scan 1 shows folding and subsequent scans do not. **c** Illustration of a mechanism by which the folded part (initially only in I_2_) gets removed by the 2nd scan when the molecule moves R-to-L, as described in the text. **d** The probability *P* (± 95% error bar) is the fraction of events that are unfolded, for the different translocation types. A total of 309 events experienced all four types in sequential order.

We propose a qualitative mechanism of the progression of unfolding during flossing in Figure 3c. Going L-to-R, motion and the field force at pore 2 are aligned, which promotes folds eventually moving through pore 2 and into channel 2. Subsequently, R-to-L motion pulls only the region of DNA that is under tension via Tug-of-War back through pore 2, despite the counter field force at pore 2, while folds not under Tug-of-War tension experience only the field force and thus remain in channel 2 and away from pore 2. Figure 3d shows the ratio of unfolded events for the progression of translocation types that each of the 309 events went through. Thus the statistics in each column are from exactly the same group of molecules, and are the same data as Figure 2. The 35% unfolded probability through the initial single pore capture (pore 1 in our device) is consistent with other single pore studies. ^1,9,33^ Following co-capture, 66% of 1st scans are unfolded, which is consistent with tug-of-war data without flossing.^28^ By the 2nd scan, only 6 molecules out of 309 (2%) remain folded, and only 5 remain folded by their last scan. Thus the flossing process effectively linearizes the DNA molecules through the nanopore sensors at high probability.

### Tag Data Analysis: A Single Scan View

The multi-scan data set (constituting L-to-R and R-to-L scans for each pore) contains rich information regarding the underlying tag binding profile and translocation physics. Each individual scan taken from the two pores provides a snapshot of the translocation process for the portion of DNA being scanned. There are two ways to assess the translocation velocity from the tag. The first approach is to quantify the speed of a tag as moves through a single-pore (‘dwell time estimation’), which is based on dividing the tag blockade duration (width-at-half-maximum) into the membrane thickness (35 nm). The second approach is unique to dual nanopore technology, and is to assess tag speed as it moves from the first pore (Entry) to the second pore (Exit). This entry-to-exit time is in reference to the time the tag resides in the common chamber above both pores, and is also referred to as the pore-to-pore time. In pore-to-pore speed estimation, the pore-to-pore time is divided into the measured distance between the pores for each chip (Table S1). By definition, the pore-to-pore approach utilizes correlation between the two current signals I_1_ and I_2_, since the time starts when the tag leaves the Entry pore and ends with the tag enters the Exit pore. We note that, at the voltages applied, DNA in the common reservoir is expected to be fully stretched between the pores^28^ (0.34 nm/bp).

Figure 4 shows an example of an adjacent pair of scans in a multi-scan event, and demonstrates how inter-tag separation distances can be estimated from each scan. For the L-to-R scan (Figure 4a,c), tag A then tag B move through pore 1, and about 1 ms later they move through pore 2. The signal pattern reveals that tags A and B are spatially closer than the distance between the pores (0.64 *µ*m, chip E, Table S1). The FPGA is monitoring I_2_ for two tags during the control logic. During the waiting period after detecting A and B in I_2_, a third tag C passes through pore 1. Visually, it is also clear from the pore-to-pore transit times of A and B that tags B and C are roughly two times farther apart than the distance between the pores. Upon changing V_1_ to promote motion in the reverse direction R-to-L (Figure 4b,d), the first observable tag in I_1_ is C passing back through pore 1. Again, the logic detects A and B in I_2_, and this time both tags pass through pore 1 before the logic triggers the V_1_ to promote L-to-R motion. Seeing three tags within I_1_ was a result of three physical tag separations that are close enough to accommodate 3-tag pore 1 transit within the 2-tag pore 2 detection time window of I_2_, as implemented on the FPGA. More common was to see two tags reliably in both pores, as detailed in the next section.

**Figure 4:**
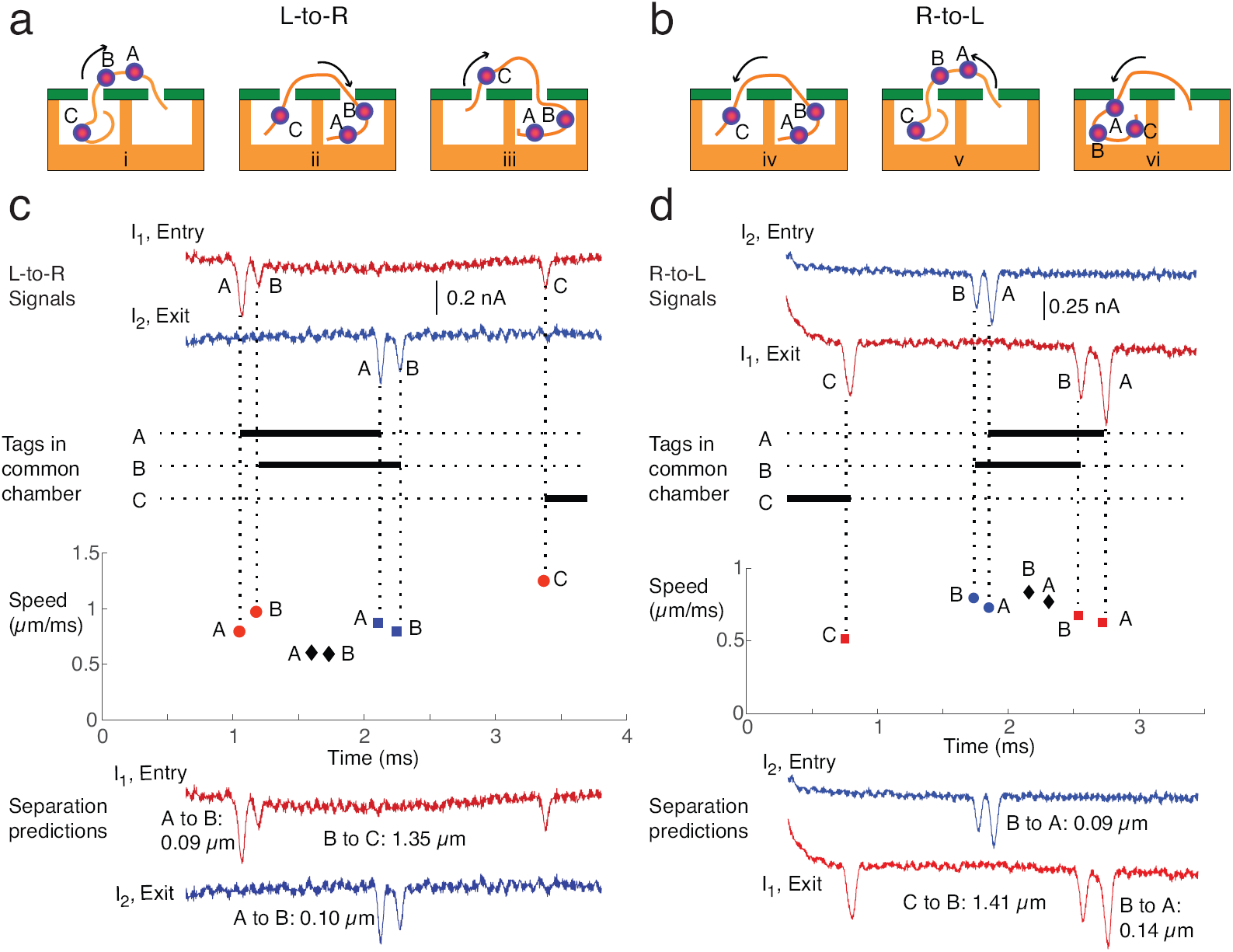
Estimating inter-tag separation distances from dual current signals generated during a multi-scan experiment. The **a** L-to-R and **b** R-to-L illustrations help visualize the relative tag locations that are revealed by the scan signals. The **c** L-to-R and **d** R-to-L signals were from adjacent scans of a co-captured molecule that was scanned for 48 cycles. In L-to-R, pore 1 is the Entry pore for a tag while pore 2 is the Exit pore. In R-to-L, pore 2 is the Entry pore for a tag while pore 1 is the Exit pore. Entry and Exit are thus relative to the direction of motion of a tag as it passes from pore to pore. The signals and inferred number of tags in the common chamber between the pores versus time are plotted. Illustration (ai) visualizes the period when A and B are in the common chamber, while (aii) visualizes the period after B exits but before C enters the common chamber, etc. The Speed plots shows the computed tag speeds at the Entry and Exit pores, based on tag duration divided into membrane thickness, and tag pore-to-pore speeds computed as the known distance between the pores divided by the pore-to-pore time. Inter-tag separation distance predictions are computed by multiplying the mean pore-to-pore speed within a scan by the time between detected tag pairs, and adding the membrane thickness as a correction (main text). The voltages were set to V_1_ = 250 mV for L-to-R and 600 mV for R-to-L, with V_2_ = 400 mV held constant (Chip E, Table S1). The FPGA monitored I_2_ for *N* = 2 tags (exit signal L-to-R, entry signal R-to-L), though 3 tags were visible in I_1_ in both directions.

For an L-to-R scan, a tag blockade in I_2_ corresponds to a tag exiting the common reservoir, making pore 2 the Exit in L-to-R, whereas moving R-to-L means a tag in I_2_ is entering the common reservoir. While I_2_ shows two tags, the third tag present in I_1_ for both scan directions yields an opportunity to quantify two tag-pair separation distances. For the signals shown, the number of tags between the pores are plotted, and the tag speeds based on dwell-time and pore-to-pore speeds are also plotted. These speed values versus time provides a glimpse into how the molecule is moving during co-capture tug-of-war, along with the illustrations added for visualization (Figure 4a,b). The pore-to-pore speed is modestly faster for an R-to-L direction (0.8 vs. 0.6 *µ*m/ms), which is consistent with a larger voltage differential for R-to-L motion (V_1_ = 600 mV, V_2_ = 400 mV) than for L-to-R motion (V_1_ = 250 mV, V_2_ = 400 mV).

We estimate tag-to-tag separation distances (Figure 4c,d bottom) by multiplying the mean pore-to-pore speed within a scan by the tag-to-tag times recorded within that scan, and adding the membrane thickness. Membrane thickness is added to account for the added spatial separation that is equivalent to either tag passing the length of a pore, since tag-to-tag times are computed from the rising edge of a detected tag blockade to the falling edge of the next detected tag blockade. The tag-to-tag distance estimates are shown for both the L-to-R scan and the R-to-L scan. The results suggest that each separation prediction will vary across the two different scan directions. Tag-pair separation distance predictions will also vary due to differences between the tag pore-to-pore speed and the true speed profile during the tag-to-tag time for a given pore. That is, the assumption that the speed between the tags is constant and equal to the pore-to-pore speed is not exactly correct. It likely that this assumption is better when the tag-to-tag times are shorter than the pore-to-pore times, which is the case for tags A and B but not tags B and C in Figure 4. In any case, if the assumption that the DNA is traveling at the constant pore-to-pore speed is true on average, we would expect the average of many re-scans to predict accurate separation distances between any two sequential tags. We next compute the average of multiple scan predictions and test how well the predictions line up with the known separations from the model tagged-DNA reagent.

### Combining Scans to Improve Tag-to-Tag Distance Predictions

We examined the error-reduction performance of averaging the distances obtained from individual scans within a multi-scan event. Five different multi-scan events with at least 30 cycles are reported in Table 1. For each event, the averaged distance estimates are shown for each scan direction, and using pore 2 estimates alone as well as merging the pore 1 and pore 2 estimates. The table reports the number of cycles, which is equal to the number of scans in each direction, and the number of tag-pairs that contributed to each separation distance estimate. In all cases, there are fewer distance estimates than scans. For example, for event (v) that had 65 scans in each direction, and for the R-to-L direction, 57 scans produced detectable tag pairs in I_2_ while 62 tag pairs were detected in I_1_ for a total of 119 separation estimates. The attrition is because the probability of a missing a tag within any one scan increases with cycle count, as described by equation (1).

**Table 1:**
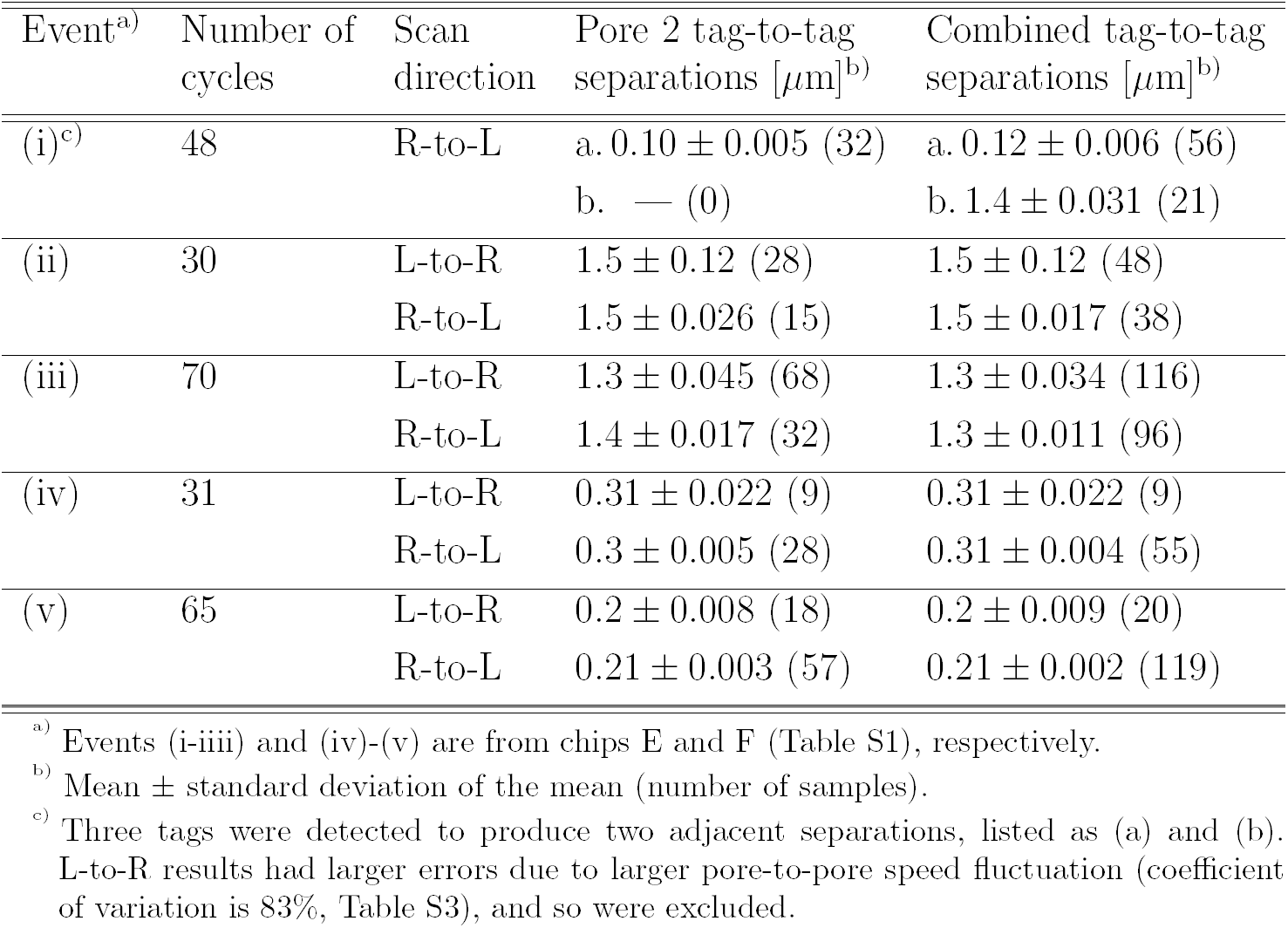
Summary of separation statistics from five multi-scan events.

| Event<sup>a)</sup> | Number of cycles | Scan direction | Pore 2 tag-to-tag separations [$\mu\text{m}$]<sup>b)</sup> | Combined tag-to-tag separations [$\mu\text{m}$]<sup>b)</sup> |
| --- | --- | --- | --- | --- |
| (i)<sup>c)</sup> | 48 | R-to-L | a. $0.10 \pm 0.005$ (32)<br>b. — (0) | a. $0.12 \pm 0.006$ (56)<br>b. $1.4 \pm 0.031$ (21) |
| (ii) | 30 | L-to-R | $1.5 \pm 0.12$ (28) | $1.5 \pm 0.12$ (48) |
| | | R-to-L | $1.5 \pm 0.026$ (15) | $1.5 \pm 0.017$ (38) |
| (iii) | 70 | L-to-R | $1.3 \pm 0.045$ (68) | $1.3 \pm 0.034$ (116) |
| | | R-to-L | $1.4 \pm 0.017$ (32) | $1.3 \pm 0.011$ (96) |
| (iv) | 31 | L-to-R | $0.31 \pm 0.022$ (9) | $0.31 \pm 0.022$ (9) |
| | | R-to-L | $0.3 \pm 0.005$ (28) | $0.31 \pm 0.004$ (55) |
| (v) | 65 | L-to-R | $0.2 \pm 0.008$ (18) | $0.2 \pm 0.009$ (20) |
| | | R-to-L | $0.21 \pm 0.003$ (57) | $0.21 \pm 0.002$ (119) |

<sup>a)</sup> Events (i-iii) and (iv)-(v) are from chips E and F (Table S1), respectively.
<sup>b)</sup> Mean ± standard deviation of the mean (number of samples).
<sup>c)</sup> Three tags were detected to produce two adjacent separations, listed as (a) and (b). L-to-R results had larger errors due to larger pore-to-pore speed fluctuation (coefficient of variation is 83%, Table S3), and so were excluded.

We can assess the correspondence between the averaged separation distances in Table 1 and the known inter-tag separations that are possible from the position map (Figure S6). Known distances are computed using the conversion 0.34 nm/bp, which assumes the DNA in the common reservoir is fully stretched between the pores. For event (i), only the R-to-L scan directions were combined and reported for event (i), since the L-to-R data showed significant variation in the pore-to-pore speed (described in SI Section 11). Note that the two scans shown in Figure 4 are from event (i), which pathologically generated two tag-pair estimates in pore 1 current for the reason described in that figure. If we assume tag 6 is absent for the molecule and that tags 4, 5 and 7 are present, the adjacent tag-pair separations for event (i) have their closest match among all possible adjacent tag-pair permutations that are possible according to the position map. Specifically, the map shows 0.1 and 1.5 *µ*m adjacent distance between tags 4-5 and 5-7. It is reasonable to assume that a tag (i.e., tag 6) is absent, as discussed previously (Figure S2e).

Events (ii-v) in Table 1 show very consistent results across both scan directions, and between both pores when comparing Pore 2 results with Combined results. For these events, only a single separation distance estimate was produced, which is most common for control logic that uses *N* = 2 tag detection in I_2_ to trigger direction switching. In terms of comparing averaged separation distances and the known inter-tag separations, events (ii) and (iii) correspond most closely to 1.5 *µ*m and 1.3 *µ*m distances between 5-7 and 6-7, respectively, with the 1.5 *µ*m value possible if the tag 6 position is assumed vacant for the molecule of event (ii). And events (iv) and (v) correspond most closely to 0.3 *µ*m and 0.2 *µ*m distances between 4-6 and 5-6, respectively, with the 0.3 *µ*m value possible if the tag 5 position is assumed vacant for the molecule of event (iv).

Consistent with the thesis of this work, the correspondence between distance estimates and map-possible permutations is generally not as clean for the individual scans (e.g., Fig. 4) as it is for the averaged scans, and is impossible for scans where tags are missed (representative examples are shown in Figs. S11, S13 and S14). This demonstrates the value of error reduction by averaging across a multi-scan data set generated for each molecule. Additional data on the velocity profiles for events (i-v) in Table 1 and data for four additional multi-scan events are reported in Table S3. The error on each separation estimate is obtained as the error on the mean over the group of estimates, with significant reduction of error achieved through averaging.

The data in Table 1 show the power of the flossing approach when the events have tag-to-tag times in I_2_ that are unambiguously attributable to the same physical set of tags (representative scans with both I_1_ and I_2_ signals are provided in Figures S7-S14). In other data, however, when a tag is missed in I_2_ within a scan, the two-tag scanning logic will eject the molecule or subsequently shift to a new physical tag pairing on the same molecule, which creates a register-shift in the tag-to-tag time data. An example of this is event (vi) in Table S3 with the register-shift scan signals shown in Fig. S11. While this complexity can be visually observed in the data and accommodated manually, we next sought to develop an alternative approach that could detect and automate analysis for such register shifts.

The alternative method presented next is based on aligning the signal in the time domain based on tag blockade proximity, with the aim of automating the binning of tag-pair times, particularly where there is greater ambiguity in assigning such times across scans. Our own prior work has shown that time-based signal alignment of nanopore data can increase the value of multiple nanopore reads in the context of sequencing through homopolymer regions.^34^ The time-based signal alignment strategy is described below, as applied to two multi-scan events shown in Figure 5. The first in Figure 5a provides an example where tag-pair matching from scan to scan is not ambiguous, while Figure 5b provides an example where tag-pair matching from scan to scan is ambiguous due to the aforementioned register-shifting effect (Figure 5a,c events are events (vii),(vi) in Table S3, respectively).

**Figure 5:**
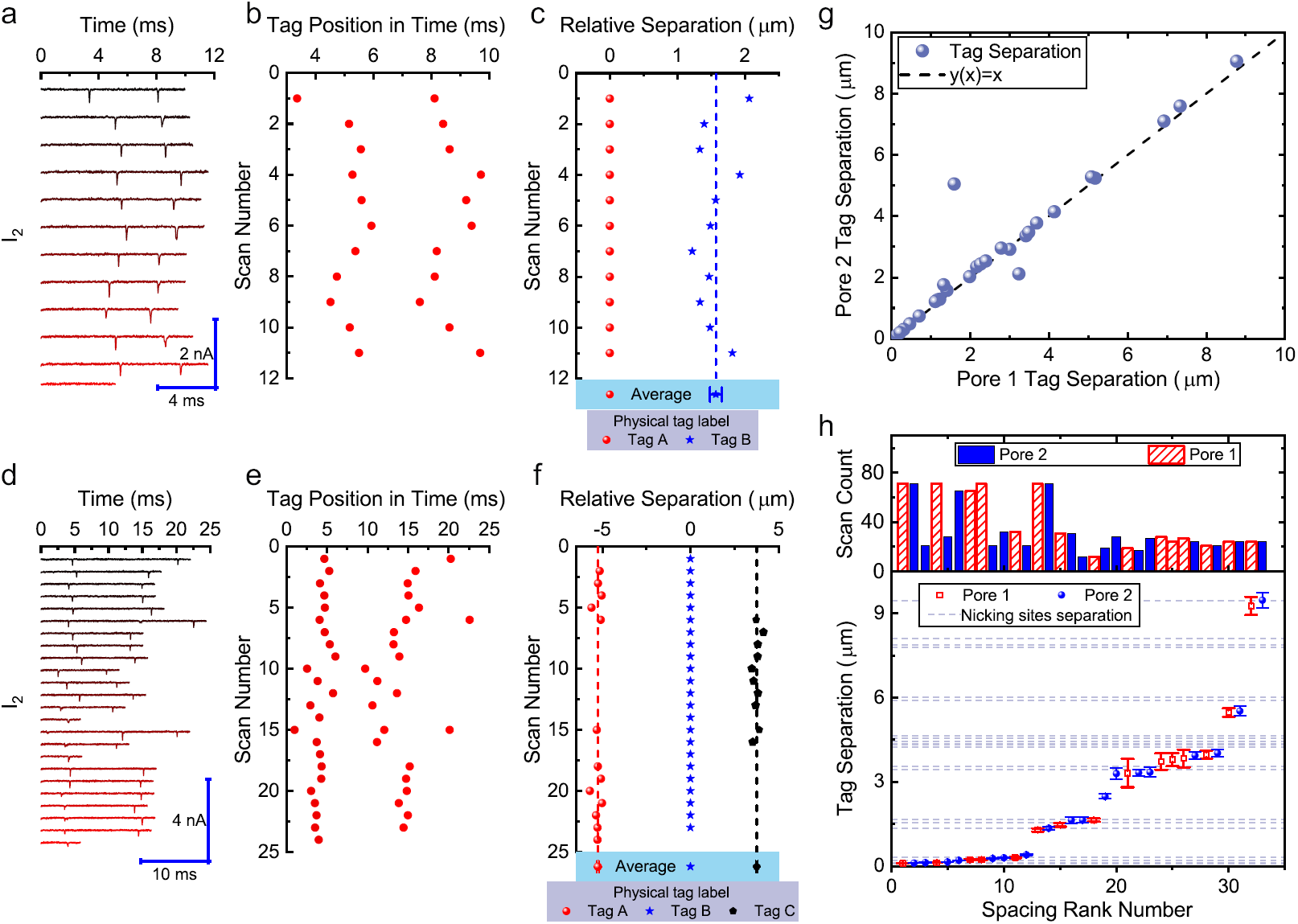
Tag position rescaling and tag spacing distribution using time-based signal aligment. **a** Raw L-to-R scans from pore 2 for a multi-scan event with control logic triggered to probe two tagged sites. The time-axis is plotted relative to the beginning of each scan. **b** Temporal positions for each tag obtained from **a. c** Aligned relative tag separations from **b** with resultant average separation distances between denoted tags A-B. **d** Raw L-to-R scans from pore 2 for a multi-scan event with control logic triggered to probe two tagged sites, but produced data covering three tagged sites due to an intermediate missed tag at scans 6 and 15 (offline analysis detects all 3 tags even where online control does not). **e** Temporal positions for each tag obtained from **d. f** Aligned relative tag separations from **e** with resultant average separation distances between denoted tags A-B and B-C. **g** Averaged separation distances between tags obtained from pore 2 versus pore 1. The linear correlation of the data indicates strong measurement reproducibility between the pores. **h** Calibrated tag separations arranged in ascending order using pore 1 and pore 2 estimates, with tag relative separation plotted versus separation ranking by size (spacing rank number). The plateaus are close to the expected permutations of tag-pair separations that are possible for the model MS-tagged *λ*-DNA reagent (horizontal lines, Figure S6). Tag separations with at least 10 scans were included, with the histogram above shows the scan count per data point. **a**-**f** data are from chip A and **g**-**h** data are from chips A, E and F (Table S1).

In the time-bases signal alignment method, the temporal position of each tag relative to the starting time of each scan is first computed. To facilitate alignment, the method must tolerate potentially large differences in tag event shape, and so the tag analysis procedure was modified and based on fitting a model function to each peak based on the convolution of a box with a Gaussian function (SI Section 10). This model can characterize tag blockades that are both broad/rectangular in character or narrow/Gaussian-like (see Figure S15 for examples of fitted scans). The extracted tag positions in time are plotted versus scan number, in Figure 5b,e. The scan to scan signals show a relative translational offset, arising from the fact that different portions of the molecule are observed in each scan. There is also stochastic variation in molecular motion, arising from Brownian fluctuations. These effects complicate correct association of tags across multiple scans, that is ensuring that the tags observed in a given scan correspond to tags at the same sequence position in adjacent scans for the same scan direction.

In order to align scans in a systematic way, the algorithm automatically groups tags and removes the translational offset across the scans. Figures 5c,f show examples of aligned events and SI section 10 provides a detailed description of the approach. Essentially, the algorithm works by assuming that at least two tags are shared between two successive scans. In order to identify one of the shared or “common” tags, the algorithm brings each potential tag pair in the two scans into alignment by shifting one scan relative to the other. Note that only translational offsets are applied, i.e., there is no overall dilation of the time-scale. For each one of these possible test alignments, the algorithm computes a measure of alignment error based on the summed squared difference between distinct tag pairs in the test alignment. The algorithm identifies the pairing that yields minimum error as true common tags and implements the translational shift that brings this pair into alignment. Note that this approach yields both the translational offset between the scans and a correspondence table of shared tags between the scans. Working iteratively across all scans in a set the tags observed across the scan can be grouped together and translational offsets removed. The tag group with the largest set of scans is defined to be the origin tag and each scan is shifted so this origin is set to zero. This algorithm outputs a final barcode, or set of averaged relative tag positions, for each set of single molecule scans. The final barcode estimates for the two multi-scan events considered are shown in Figure 5c,f, showing values that are very close to those reported using the method of combining distances presented above (see events (vii),(vi) in Table S3). The error on a tag position in the barcode is obtained as the error on the mean over the group of scans associated with the tag, with significant reduction of error achieved through averaging of positional information over multiple scans.

The outputted barcodes are in units of time. In order to calibrate the scans to units of distance, we first use an aggregate translocation velocity corresponding to the scan set. The aggregate translocation velocity is computed as the mean of a subset of the pore-to-pore speeds measured within a multi-scan event, using only those speeds for which the scan displayed a conserved number of tags in both pores. The barcodes are then calibrated in units of distance by multiplying by the mean pore-to-pore velocity. The reported error on the final calibrated tag positions incorporates the error on the velocity and the error in separation via standard propagation. The set of possible tag separations extracted from these barcodes shows good reproducibility between the pores (Fig. 5g) and correspondence with the expected separations from the *λ*-DNA tag map (Fig. 5h). Note that the sharpness of the plateaus indicate the precision achieved through averaging of multiple scans. In particular, when we sort the extracted tag separations by size, we find that the data shows distinct plateaus that correspond to the expected separations in the map. Separations that fall off the expected spacings could arise from non-specific binding (e.g., tags attached at random nicks present non-specifically in *λ*-DNA), or offsets caused by imperfections in the tag blockage analysis algorithm. Figure 5g,h shows data from the L-to-R portion of 33 separate multi-scan estimates with an average velocity of *v* = 0.56 *±* 0.04 *µ*m/ms and a minimum of 10 scans per tag pair. We also plotted the scan count aligned with each data of the 33 estimates, with higher scan counts reducing the variance of each data point.

The data in Figures 4 and 5h show that the closest spacing available with the model reagent (300 bp or 100 nm) is well resolved. In single-pore work, sub-4 nm diameter membrane-based pores that hug PNA-based motifs (15 bp footprint) have demonstrated 100 bp inter-tag distance resolution,^35^ while pipette-based pores have resolved at low as 200 bp. ^6^ We did not explore the lower limit of inter-tag distance resolution here, but observe that since our peaks are well resolved for the 300 bp separation (e.g., visible space between tags in Figure 4 traces), and since slower passage and higher bandwidth are knobs that can be turned in our setup, we anticipate a lower limit below 300 bp should be achievable. It should be possible to resolve tags that are spaced modestly farther apart than the membrane thickness, in principle (∼ 105 bp for the current chips). This can be examined in future work with different model reagents.

## Conclusion

We have developed an approach that first traps and linearizes an individual, long DNA molecule in a dual nanopore device, and then provides multi-read coverage data using automated “flossing” control logic that moves the molecule back-and-forth during dual nanopore current sensing. From the point-of-view of dual pore technology development, our rescanning approach overcomes a key challenge: while maximally long trapping-times can be achieved by balancing the competing forces at each pore, sub-diffusive dynamics will persist as speed is reduced,^28^ undermining mapping-based approaches that rely on a correspondence between the time at which tags are detected and their physical position along a DNA. In our approach sub-diffusive dynamics are avoided during bidirectional scanning of the molecule by using speeds that permit reliable tag detection while being high enough to avoid broad translocation time distributions. By enhancing the quality of information that can be extracted from a given DNA carrier strand, we feel this is a step towards addressing a key challenge in single-molecule technology: genome-scaling. Genome scaling is the potential to move beyond experiments with short DNA constructs and tackle complex, heterogeneous samples containing fragments in the mega-base size drawn from Gbp scale genomes. For genome scaling to be feasible, single-molecule reads must have sufficient quality (e.g. contain sufficiently low systematic and random errors) to enable alignment to reference genomes and construct contigs from overlapping reads drawn from a shared genomic region. This is the only way to identify long range structural variations that are masked by short read methods (i.e., via NGS), and in some cases masked even by long-read sequencing. ^36^ Future work will include developing a new model reagent with improved tagging efficiency, and optimizing the multi-scanning logic for increased scan number. We also plan to apply the approach to longer molecules with more complex tagging patterns, possibly using repetitive scanning at targeted regions to gradually explore the barcode structure. In the applied context of epigenetics, our technique can combine sequence-specific label mapping, using the same chemistry here or other nanopore-compatible schemes that have a low spatial footprint per label, ^13,37^ with methylation-specific label detection.^14^ This would help meet the need for technologies that perform long-range methylation analysis.^38^

## Experimental Section

### Preparation of mono-streptavidin tagged *λ*DNA reagent

5 *µ*g of commercially prepared *λ*DNA (New England Biolabs) was incubated with 0.025 U of Nt.BbvC1 in a final volume of 100 *µ*l of 1 X CutSmart buffer (New England Biolabs) to introduce sequence specific nicks in *λ*DNA. The nicking reaction was incubated at 37^*o*^C for 30 minutes. The nicking endonuclease Nt.BbvC1 has the recognition sequence, 5’-CC*↓*TCAGC -3’ and there are 7 Nt.BbvC1 sites in *λ*DNA. Nick translation was initiated by the addition of 5 *µ*l of 250 *µ*M biotin-11-dUTP (ThermoFisher Scientific) and 1.5 U of E.coli DNA polymerase (New England Biolabs) and incubated for a further 20 minutes at 37*°*C. The reaction was quenched by the addition of 3 *µ*l of 0.5M EDTA. Unincorporated biotin-11-dUTP was removed by Sephadex G-75 spin column filtration. To create mono-streptavidin tagged *λ*DNA complex, mono-streptavidin was added to the G-75 purified biotin labeled DNA to a final concentration of 50 nM and incubated at room temperature for 5 minutes to allow the biotin - mono-streptavidin interaction to saturate. The mono-streptavidin tagged *λ*DNA complex was then used directly in nanopore experiments.

### Fabrication process of the two pore chip

We described the fabrication protocol in our previous work. ^33^ Briefly, we prepared the microchannel on glass substrate and SiN membrane on Si substrate separately. The all-insulate architecture minimized the system capacitance. Thus the noise performance is optimized. We initially dry-etch two “V” shape, 1.5 *µ*m-deep micro-channel on the glass in a 8 mm *×* 8 mm die, with the tip of the “V” 0.4 *µ*m away from each other. Considering we can not grow SiN membranes on glass, we then deposit 400 nm-thick LPCVD SiN, 100 nm-thick PECVD SiO2, and 30 nm-thick LPCVD SiN on Si substrate. To transfer the 3-layers film stack to glass substrate from the Si substrate, we anodic-bonded the two substrates, with the micro-channel on glass facing the 3-layers film stack on Si. To remove the Si substrate, we first dry-etched away the 430 nm SiN on the backside of Si. Then we etched away the Si substrate using hot KOH, revealing the 3-layers films stack on the glass. The 3-layers films stack provides mechanical support to cover the micro-channel on glass, while it is too thick for nanopore sensing. So we had to open a window in the center for nanopore. To achieve that, we dry-etched a 10 *µ*m x 10 *µ*m window in the center through the 400 nm-thick SiN mask into the 100 nm-thick SiO2 buffer layer. Then we etched away the leftover 100 nm-thick SiO2 layer using hydrofluoric acid, revealing the single 35 nm-thick SiN membrane layer. At last, we drilled two nanopores through the membrane using Focused Ion Beam at the tip of the two “V” shaped channels.

### Nanopore experiments

We performed all the nanopore experiments at 2 M LiCl, 10 mM Tris, 1 mM EDTA, pH=8.8 buffer. The two pore chip was assembled in home-made micro-fluidic chunk, which guide the buffer to channel 1, channel 2, and the center common chamber. Ag/AgCl electrodes were inserted to the buffer to apply voltage and measure current. The current and voltage signal was collected by Molecular Device Multi-Clamp 700B, and was digitized by Axon Digidata 1550. The signal is sampled at 250 kHz and filtered at 10 kHz. The tag-sensing and voltage control module was built on National Instruments Field Programmable Gate Array (FPGA) PCIe-7851R and control logic was developed and run on the FPGA through LabView.

### Data analysis

All data processing was performed using custom code written in Matlab (2018, MathWorks). The start and end of each scan and event are extracted from the FPGA state signal (SI) for offline analysis. During real-time tag detection on the FPGA, the presence of tag is detected in the control logic if any sample falls 70 pA below the baseline and lasted at least 12 *µ*s. During off-line analysis, for the analysis reported in Figure 4 and Table 1, tag blockade quantification during each scan is performed as follows: the open pore baseline standard deviation is calculated using 500 *µ*s of event-free samples (*σ*); the DNA co-capture baseline *I*_DNA_ is determined using the mean of 100 tag blockade-free samples; a tag blockade candidate is detected where at least one sample falls below *I*_DNA_ − 6*σ*, i.e., sufficiently below the DNA co-capture baseline; a tag blockade is quantitated where the blockade candidate has samples that return within 1*σ* below *I*_DNA_, and the tag duration is computed as the full width at half minimum (FWHM), where the half minimum is halfway between the lowest sample below the DNA baseline and the DNA baseline. The alternative tag profile characterization via least-squares fitting that was utilized for the alignment strategy data in Figure 5 is described in SI section 12. Tag-to-tag times are computed from rising edge to falling edges using the FWHM time transition (edge) points, and pore-to-pore times use the rising edge of the tag blockade at the entry pore, to the falling edge of the corresponding tag blockade at the exit pore. Pore-to-pore times were computed by assigning entry tags to have one matching exit tag, utilizing the first exit tag not previously assigned and within a time limit of 10 ms. Cases where missed tags in analysis produced incorrectly assigned pore-to-pore times occurred ∼9% of the time (see tag-pair and pore-to-pore time counts in Table S3), and were manually trimmed. Pore-to-pore times were utilized to compute pore-to-pore speed on a per scan basis (Figure 4, Table 1, Table S3) or in the aggregate with the alignment strategy data (Figure 5) as described in the main text and SI section 12. Compensation of transient decay in I_1_ following step changes in V_1_ is described in SI section 12.

## Supporting Information

Supporting Information is available from the Wiley Online Library or from the author.

## Acknowledgement

This work done in Santa Cruz was performed and financially supported by Ontera, Inc. The work done at McGill University was financially supported by the Natural Sciences and Engineering Research Council of Canada (NSERC) Discovery Grants Program (Grant No. RGPIN 386212).

