## Supplemental for "Flossing DNA in a Dual Nanopore Device"

### Flossing DNA in a Dual Nanopore Chip Supplementary Material

**Images of 2 Pore Chips and Applied Voltage** 2

**Tug-of-war Experiment with DNA** 4

**Tug-of-war Experiment with Tagged DNA** 6

**FPGA logic of Multi-scan Experiments** 8

**Summary of The Last Cycles in The Multi-scan Experiments** 9

**Multi-scan Experiments with Three-tags Triggering** 11

**Tag Location Map** 12

**Examples of Scans** 12

**Tag Profile Characterization via Least-squares Fitting** 16

**Tag Alignment Procedure** 18

**Tag-to-tag Separation Statistics from Nine Multi-scan Events** 22

### 1. Images of 2 Pore Chips and Applied Voltage

| Chip Name | Used in the paper | Pore to pore distance (μm) | Pore 1 diameter (nm) | Pore 2 diameter (nm) |
|---|---|---|---|---|
| A | Figure 1(b)<br>Figure 5<br>Figure S15 | 0.66 | 23 | 23 |
| | | 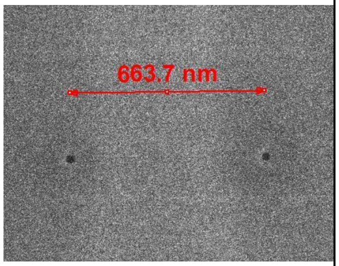 | 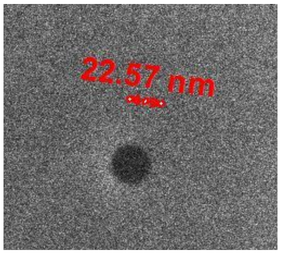 | 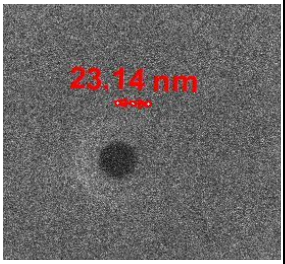 |
| B | Figure S1<br>Figure S2 | 0.67 | 27 | 29 |
| | | 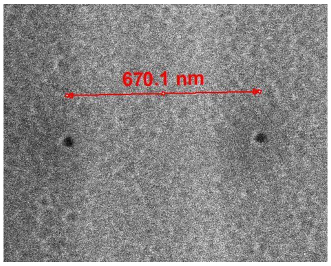 | 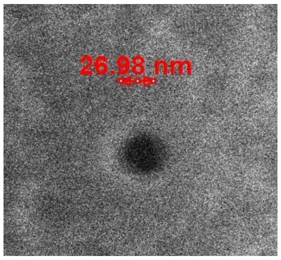 | 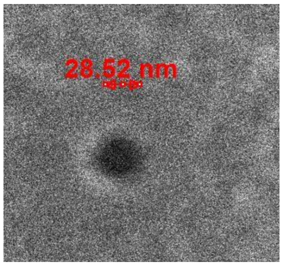 |
| C | Figure 2<br>Figure S3(a)<br>Figure S4<br>Figure 4 | 0.61 | 27 | 25 |
| | | 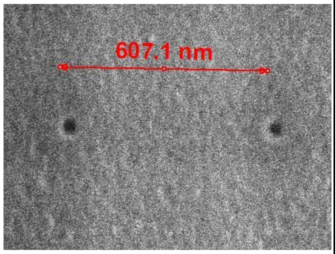 | 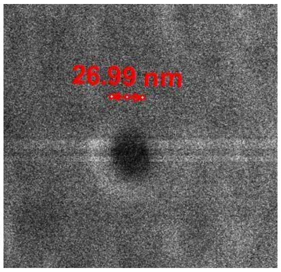 | 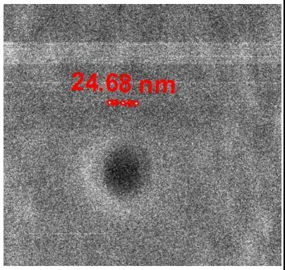 |
| D | Figure S5 | 0.65 | 29 | 29 |

| | | | | |
|---|---|---|---|---|
| | | 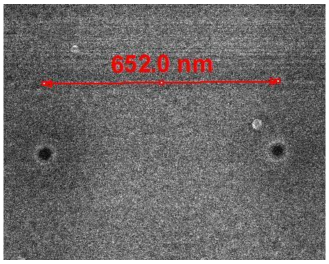  | 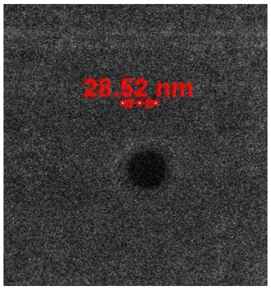  | 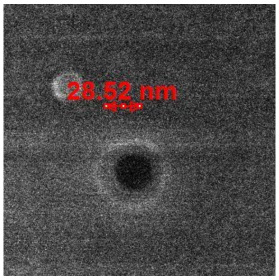  |
| E | Figure 3<br>Table 1<br>Figure 5 | 0.63 | 22 | 21 |
| | | 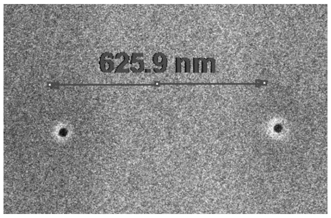  | 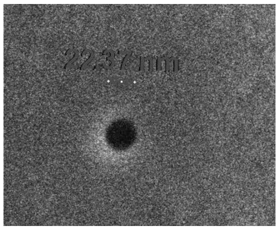 | 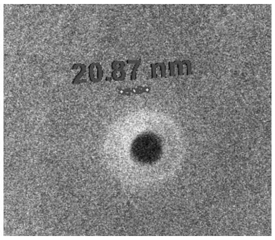  |
| F | Figure 3<br>Table 1<br>Figure 5 | 0.64 | 19 | 24 |
| | | 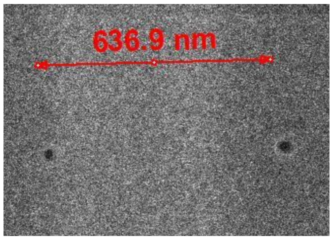  | 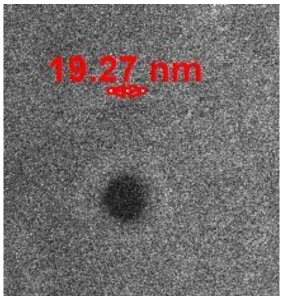  | 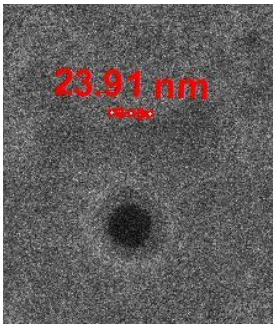  |
| G | Figure 3<br>Table 1 | 0.59 | 21 | 22 |
| | | 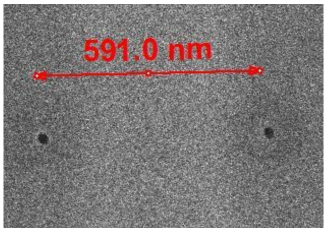  | 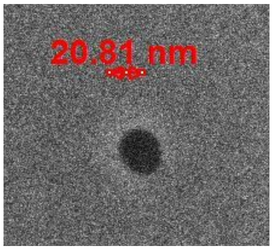  | 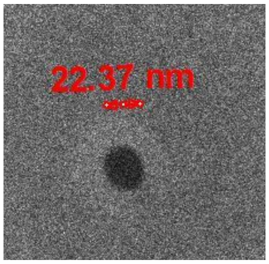  |

Table S1: The images of the chips that we used in the paper. We took these images using FEI Helios NanoLab 600i DualBeam FIB/SEM at Stanford University.

| Chip name | Applied voltage in the experiment | | | |
|---|---|---|---|---|
| | $V_{1,\ L\text{-}to\text{-}R}$ (mV) | $V_{1,\ R\text{-}to\text{-}L}$ (mV) | $V_{1,\ balance}$ (mV) | $V_2$ (mV) |
| A | 150 | 600 | 300 | 300 |
| B | 200 | N/A* | 500 | 500 |
| C | 100 | 850 | 300 | 300 |
| D | 150 | 800 | 300 | 400 |
| E | 250 | 600 | 350 | 400 |
| F | 200 | 600 | 300 | 400 |
| G | 250 | 550 | 400 | 400 |

Table S2: The voltage used in each experiment. The chip name refers to the same chip in Table S1. $V_{1,L\text{-}to\text{-}R}$ and $V_{1,R\text{-}to\text{-}L}$ refer to the low and high voltages respectively applied at pore 1 during the multi-scan experiment. $V_2$ is the voltage applied at pore 2 when the molecule is co-captured. We normally ran tug-of-war experiment using $\lambda$DNA in advance to calibrate the system. Then we assign $V_{1,balance}$ to be the voltage when the duration reaches its maximum.
* The $V_{1,R\text{-}to\text{-}L}$ of chip B is N/A because we only ran the tug-of-war experiment using this chip.

#### 2. Tug-of-war Experiment with $\lambda$DNA

Figure S1 shows the details of tug-of-war experiment with $\lambda$DNA. We always run the tug-of-war experiment with $\lambda$DNA molecules in advance to calibrate the 2-pore device. Figure S1a shows the full steps of the tug-of-war experiment with corresponding current trace $I_1$ and $I_2$ in Figure S1b. We initially filled the common chamber with 20 pM $\lambda$DNA. The idle state (state pre-i, 0-50 ms) of the system was set to be $V_1$=300 mV, $V_2$=300 mV. Once a downward spike showed up in $I_1$ at 15 ms, the FPGA measured the spike. If the spike jumps at least 70 pA below the baseline and lasts at least 0.5 ms, the system treats it as an intact molecule and get ready to progress to state pre-ii. Otherwise the system stays in state pre-i to be ready for the next trigger. That false positive spike may cause by some DNA fragments or free leftover protein when we prepare the tagged $\lambda$DNA. Then we set V1=0 mV in state pre-ii (50-70 ms) to let the $\lambda$DNA molecule relax to its equilibrium conformation. After that, we set $V_1$=-200 mV to drive the molecule back to pore 1 in state pre-iii. Notice we kept $V_1$=300 mV unchanged after the spike for 30 ms at state pre-i. The purpose was to push the molecule a distance away from the nanopore. Thus the molecule wouldn't run back that fast to show up in the exponentially decay baseline, which is hidden in the axis break around 70 ms. Because it is hard to trigger a translocation in a drifting baseline. When the molecule did come back to pore 1, it generated an upward spike around 80 ms (see the insert). Before the translocation completed, we turned off $V_1$ within 0.3 ms in state i to let the head of $\lambda$DNA dangling at the common chamber, waiting to be caught by pore 1, while keeping the rest of the $\lambda$DNA anchored in pore 1. We also increased $V_2$ to be 500 mV in state i to generate a stronger force to catch that head of the $\lambda$DNA. At around 165 ms, the head of the $\lambda$ DNA molecule reached pore 2, generating another downward spike (see the insert). We then set $V_1$ to be 400 mV to pull the other end of the molecule in the opposite direction, reaching the tug-of-war state ii. Because the pulling force in pore 1 was still weaker than that in pore 2, the molecule still slid towards pore 2. The molecule finally exit pore 1 and pore 2 sequentially at around 215 ms at state iii. Then the system went back to the idle state i for another cycle. The $I_2$ baseline did not come back to the original value in state i, which was caused by the cross-talk between the two pores[1]. $I_1$ showed a huge spike and exponentially decay baseline each time after changing its value each time, which was caused by the capacitance of the chip. An electrical diagram was shown in our previous work to explain the cross-talk and capacitance[1].

Adding the three states, pre-i, pre-ii, and pre-ii, provides two advantages. First, compared to filling the reagent in channel 1, filling the reagent in common chamber saves one step of pumping the reagent through the channel, which enables us to exchange different reagents more efficiently. Second, the single pore translocation in state pre-i screens out the short fragments. We only ran the tug-of-war experiment in state i, ii, and ii with the molecule which showed long enough duration in state pre-i, which increases the efficiency of grabbing the intact molecules.

As we stated in the main text, we define the tug-of-war duration as the time spent at state ii. We kept $V_2$=500 mV in state ii unchanged and adjust $V_1$ to measure the duration. Figure S1c shows the mean duration with errorbar. Similar to our previous work[2], the Duration (ms) Vs $V_1$ (mV) shows a bell curve. The duration reached its maximum of 110 ms at the balanced voltage $V_1$=500 mV, while the duration shows one magnitude smaller value at unbalanced voltage. Because the molecule moved faster when the net force from either pore 1 and pore 2 is larger. We also measured the single pore duration in state pre-i and plot its mean duration, 3.4 ms, in the grey area, illustrating the same molecule shows more than one magnitude shorter duration in single pore translocation compared to that in tug-of-war.

Considering the state ii is the most critical state in the process, we normally only show the current trace pair in that state. Figure S1d shows event example with $\lambda$DNA at the voltage setting of $V_1$=200 mV, $V_2$=500 mV.

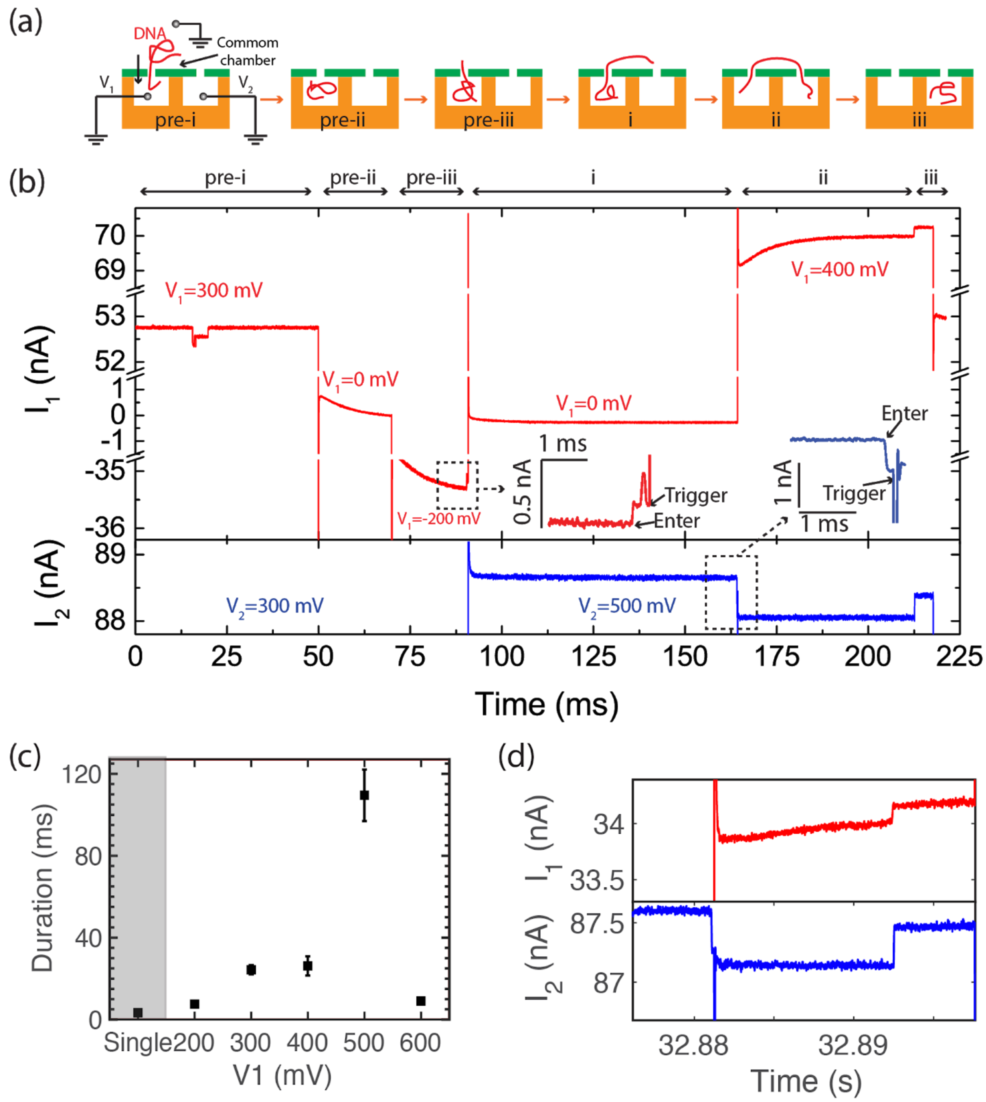


Figure S1: Full process of one tug-of-war event. (a) Illustration of each step. (b) Current trace from both pore 1 and pore 2. $I_1$ is in red while $I_2$ is in blue. We broke the y-axis of pore 1 three times to fit the different value in one figure. The insect shows the zoom-in when trigger happens. (c) The mean duration at different $V_1$ with single pore events duration. (d) Current trace pair of one tug-of-war event with $\lambda$DNA at $V_1$=200 mV.

#### 3. Tug-of-war Experiment with Tagged $\lambda$DNA

Figure S2a shows the process of constructing the mono-streptavidin bonded $\lambda$DNA complex. We described the process in our previous work[2]. Briefly, we initially nicked one strand of the $\lambda$ DNA with endonuclease enzyme Nt.BbvC1m, which nicks N=7 total sites along the $\lambda$DNA. Then we attached biotinylated dUTP at the nicking sites. At last, we conjugated the mono-streptavidin on the biotin.

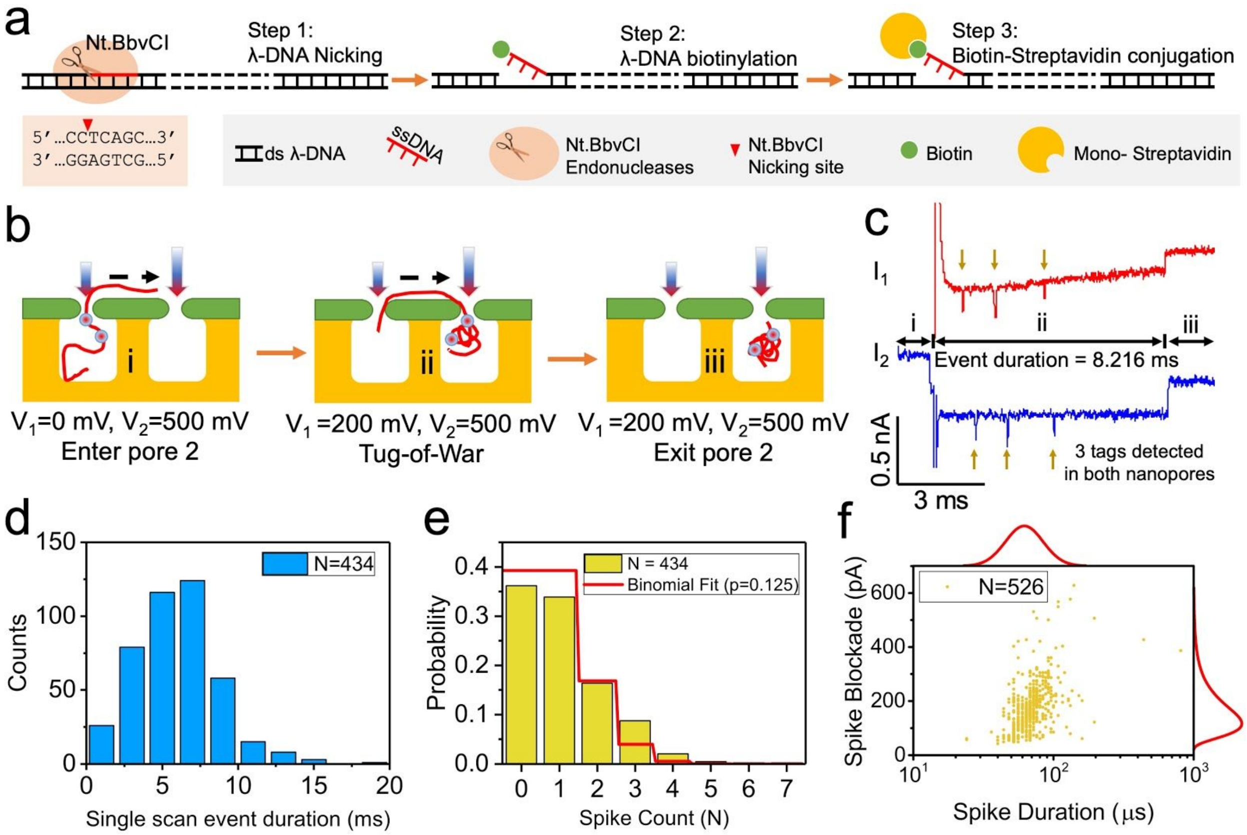


Figure S2: Tug-of-war experiment with tagged DNA. (a) Construction of a streptavidin labeled $\lambda$ DNA molecule. (b) Illustration of tug-of-war process with tagged $\lambda$DNA. (c) Current pair from one tug-of-war event. We sample the signal at 250 kHz and filter it at 10 kHz. (d) Distribution of the duration of all the tug-of-war events. (e) Distribution of the spike count in each tug-of-war event. (f) Scatter plot of Spike blockade Vs Spike Duration for $I_2$.

Figure S2b illustrates the critical steps in the tug-of-war experiment. The full process is shown in the previous section. We kept $V_2$=500 mV unchanged during tug-of-war. As soon as the head of molecule entered pore 1 at state i, we turned $V_1$ off, leaving the head of the molecule dangling in the common chamber, waiting to be caught by pore 2. Once the head of the molecule reached pore 2 at state ii, we turned $V_1$ on to be $V_1$=200 mV, reaching tug-of-war state. The molecule then kept moving towards pore 2 since force in pore 1 was weaker than that in pore 2. Ultimately, the molecule left pore 1 and pore 2 sequentially at state iii. The corresponding current from pore 1 and pore 2 is shown in Figure S2c. Label i, ii, and iii indicate the corresponding three states in Figure S2b. We defined the event duration as the time spent in state ii. We observed three spikes in both $I_1$ and $I_2$ during state ii, indicating three tags along the $\lambda$DNA. We call it a spike when the current value jumps below 3 standard deviation ($\sigma$) of the baseline and lasts at least 12 μs.

Figure S2d~f show the results of one tug-of-war experiment with mono-streptavidin tagged $\lambda$ DNA. The experiment was performed using a chip with 0.67 μm pore-to-pore distance, 27 nm pore 1 diameter, and 29 nm pore 2 diameter (Table S1).

In terms of the MS tagging efficiency on the model DNA, higher tagging efficiency naturally provides more molecules that generate multi-scan data, as described in the next section. To examine the efficiency of our model reagent, we counted the spikes in each event and show the distribution in Figure S2e. If we assume the tags have probability *p* of being bound at each DNA site, then from a binomial distribution the probability *P(k)* of detecting *k* spikes among a total or *n=7* sites within one event is given by

$$P(k) = C_n^k \, p^k (1-p)^{(n-k)}, \quad \text{where} \quad C_n^k = \frac{k!}{n!(n-k)!} \qquad \text{S1}$$

Fitting the data to equation equation S1 results in *p=0.125* (Figure S2e). At such low binding probability, we found less than 20% of the events showed more than 2 spikes. Future reagents will aim to improve on this efficiency, while the model is sufficient to generate proof-of-concept data that shows the multi-scan functionality presented in this paper. A scatter plot of the spike blockade vs.~duration characterizes the spike signal, with the distribution centering at 0.1 nA and 70 μs for the data shown (Figure S2f), providing metrics that can be used to tune the multi-scanning.

#### 4. FPGA logic of Multi-scan Experiments

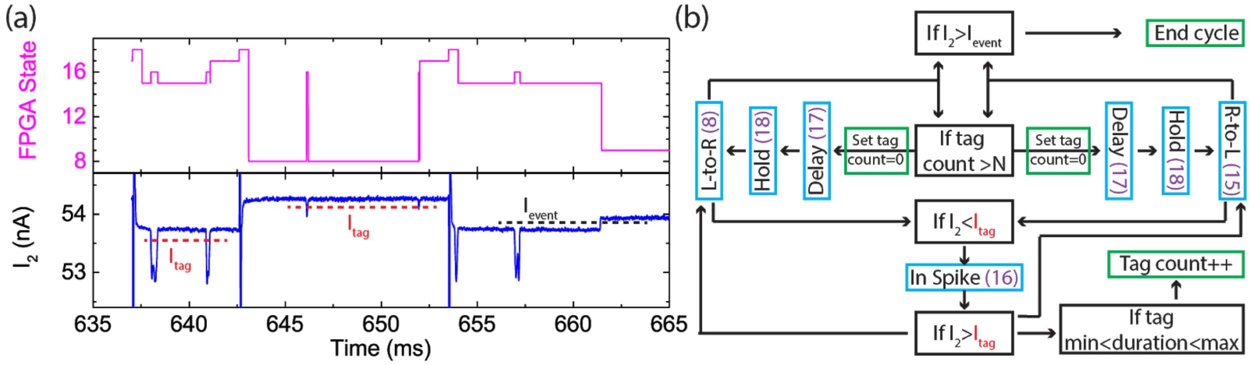


Figure S3: FPGA logic of the multi-scan experiment. (a) Signal of $I_2$ and FPGA state from the end of the event shown in figure 2. The red and black dashed line shows the triggering threshold of tag and event respectively. (b) FPGA logic flowchart. The blue boxes indicate the system state with magenta numbers refers to the FPGA state in (a). The black boxes indicate decisions. The green boxes indicate actions. The arrows between the boxes indicate how the system flows between different steps.

Figure S3 illustrates the FPGA logic of the multi-scan experiments. Figure S3a shows the signal of $I_2$ and FPGA state from the last cycle of the event in figure 2a in the main text. We hided $I_1$

here since the FPGA only triggers the spikes in $I_2$. Pore 2 performs as the sensor while pore 1 performs as the controller in our design. The FPGA State is the integer numbers reported by the FPGA to reveal its internal state. Figure S3b shows the FPGA logic flow. We designed the flow to control the system switch between two major states: L-to-R (8) when the molecule moves from left to right, and R-to-L (15) when the molecule moves from right to left. The In Spike (16), Delay (16), and Hold (18) are the transitional states. The FPGA calculate the baseline as the mean of $I_2$ up to 10 ms, while the $I_2$ samples during the In Spike (16) is excluded. And the FPGA refreshes the baseline value once the major state (8 or 15) changes. $I_{tag}$ and $I_{event}$ are relative values comparing to the baseline to trigger the spike and the end of the event. The system is in state 15 during 637 ms to 642 ms. $I_2$ jumped below $I_{tag}$ in 638 ms, triggering the system to enter state 16 when the system started to count the duration. Once $I_2$ jumped back above $I_{tag}$, the system switched back to state 15. If the tag duration is between the minimum (7μs) and maximum (2 ms) of the user settings, the internal tag count increased by 1. Similar process happened in 641 ms, when the tag count increased by another 1. Once the tag count increased beyond a user input N, which we set N=2 in this experiment, the system entered the process of switching to the other major state (8). To avoid the spike showing up too soon after state switching, we designed a delay state 17 for delaying 1.5 ms to push the last tag move further away the nanopore, which is shown in 642 ms and 652 ms. As soon as the system switch the major states, the tag counter was reset to 0, ready for the new triggers. Note there is another 0.5 ms-long hold state (18) when triggering is disabled right after switching the voltage, which is around 643ms and 654 ms in the signal. Because the system needs time to calculate the baseline value. As a result, the system missed the spike in 654 ms. Thus the tag counter did not reach N even after the two tags already showed up. As a result, $I_2$ finally jumped above $I_{event}$ at 661 ms, indicating the molecule left pore 2. Then the event ended.

### 5. Summary of The Last Cycles in The Multi-scan Experiments

Figure 2d in the main text shows the system caught the scan at the probability of p=0.89, which is pretty high. Though we still studied the last cycles to figure out the reason why the molecule escaped the multi-scan. Figure S4 shows the signal of the FPGA state and $I_2$ of the last cycles, which include the $(n-1)^{th}$ and $n^{th}$ scan.

Figure S4a shows a missing tag in the $n^{th}$ scan. The FPGA calculates the baseline at the hold (18) state. Because the baseline changed its value when the V1 changes due to cross talk. Any spikes show up during state 18 would not be detected. Missing that tag, the tag counter can not reach to the user set value N. Thus system wouldn't trigger to switch the major state to continue the multi-scan. The molecule exits the pore at 11 ms.

Figure S4b shows another case when the FPGA detected a false positive in the $(n-1)^{th}$ scam. The tiny spike around the 1.8 ms might be caused by some free protein instead of a real tag along the DNA. Thus the system switched the major state without reaching tag count N=2 at the (n-1)th state. As a result, there would be no second tag to trigger in the nth state. The molecule exits the pore at 12.5 ms.

Figure S4c shows a false negative spike in the nth scan. Sometimes the tag get stuck in the pore and produces a long spike. Once its duration exceeds the maximum value (2 ms) in the setting. The FPGA won't count it a valid one. Failing to switch to the other major state, the molecule exits at around 15 ms.

Figure S4d shows the molecule exits the pore at the delay (17) state before the FPGA switch to the other major state. The molecule exits the pore at around 5.5 ms, before the FPGA switch to the other major state at 6 ms.

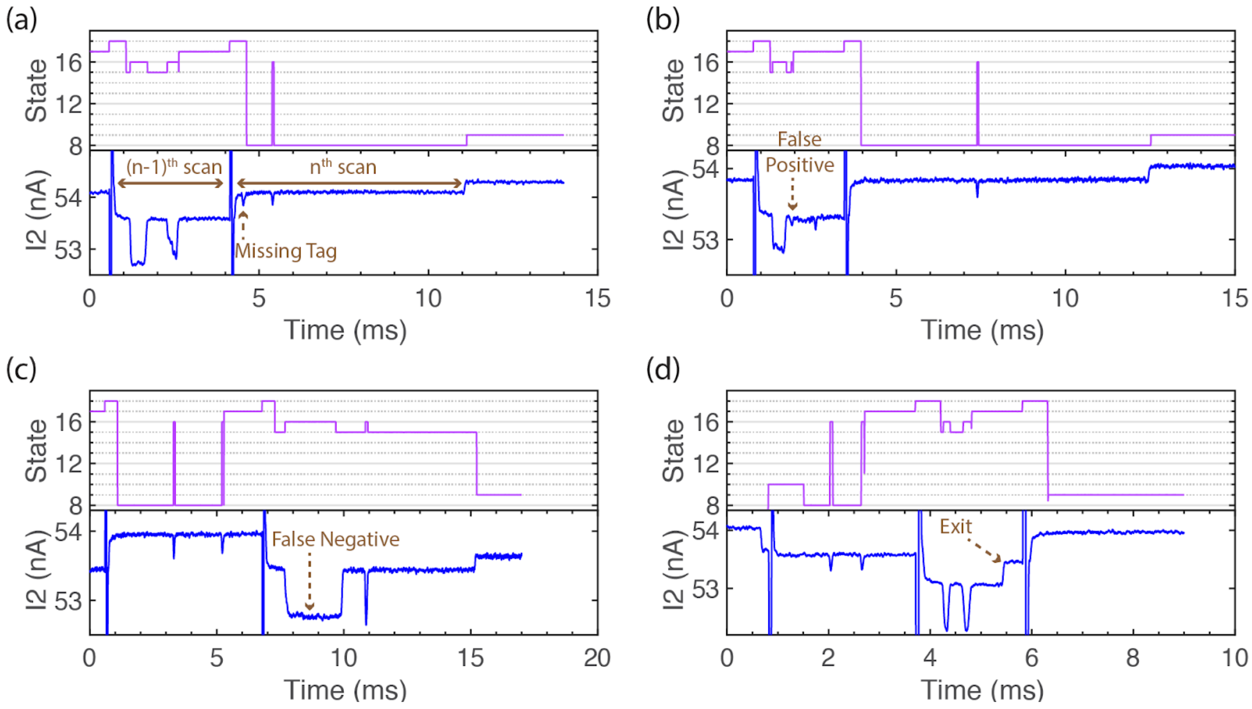


Figure S4: Four major cases that the FPGA failed to catch more scans. The plots are the signal of State from FPGA and $I_2$ in the last cycle. The definition of the state is in Figure S3b. (a) The tag shows up in the hold (18) state of the nth scan. The line with arrows mark the (n-1)th and nth scan. (b) A false positive spike in the (n-1)th scan. (c) A false negative spike in the nth scan. (d) The molecule left pore 2 in the delay (17) state of the nth scan.

We set empirical values to optimize the system to maximize the total count of scans in each event. We set spike duration with 7 µs minimum and 2 ms maximum, delay for 1.5 ms and hold for 0.5 ms. Too long delay time in state 17 would cause more failing cases in Figure S4d, while short delay time would cause more failing cases in Figure S4a. Because the tag tends to be driven back too soon if it is too close to the pore. Too long hold time in state 18 causes more failing cases in Figure S4a, while too short hold time masses up the calculation of baseline value. Wrong baseline value would cause more false positive or negative spike detection, which increases the failing cases in Figure S4b and c.

#### 6. Multi-scan Experiments with Three-tags Triggering

Figure S5 shows the experiment with three-tags triggering. Figure S5a shows the signal trace from a 6 cycle event. Figure S5b shows the zoom-in signal of the 2nd cycle. Figure S5c shows the distribution of scans count per event.

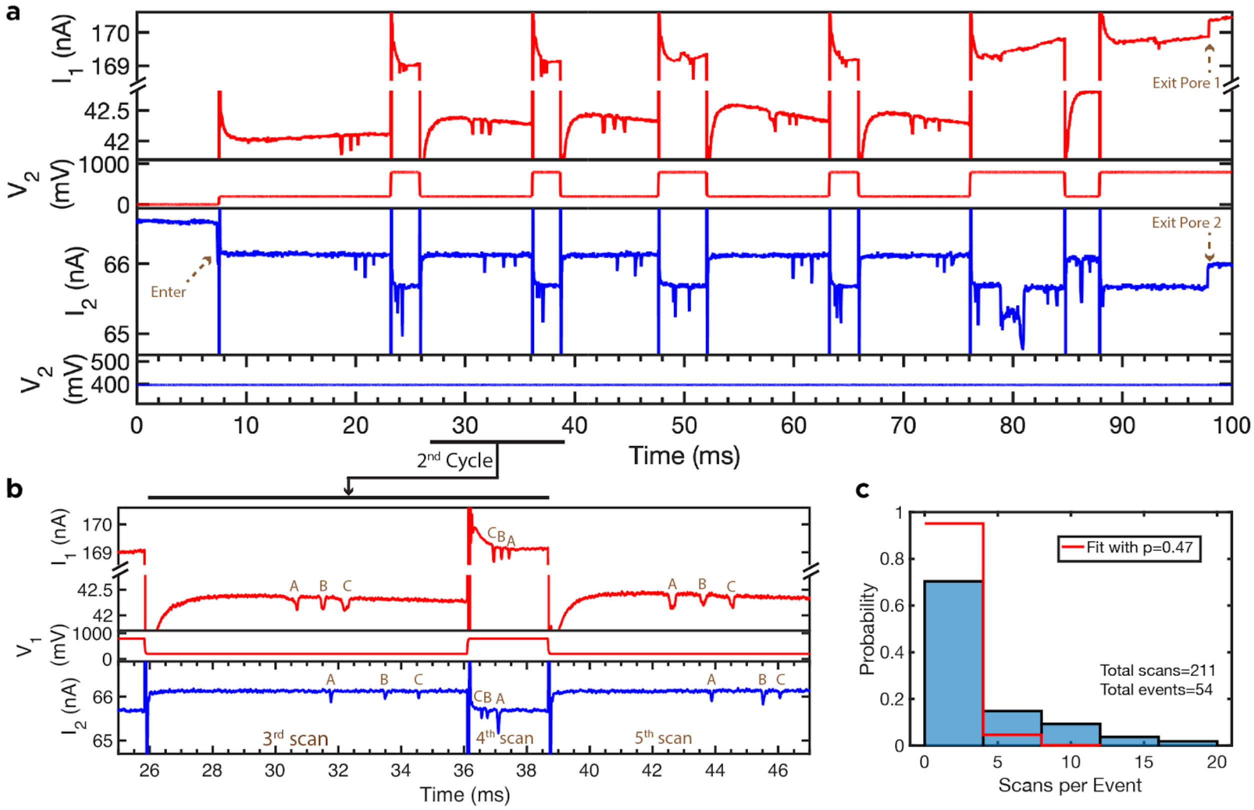


Figure S5: Multi-scan experiment with three-tags trigger. (a) Full signal trace of $I_1$, $V_1$, $I_2$, and $V_2$. We set $V_2$=400 mV during the event, $V_1$=200 mV for L-to-R scan and $V_1$=800 mV for R-to-L scan. V1 was set (b) Zoom-in plot of the 2nd cycle. (c) Distribution of the scans count per event with theoretical fitting.

#### 7. Tag Location Map

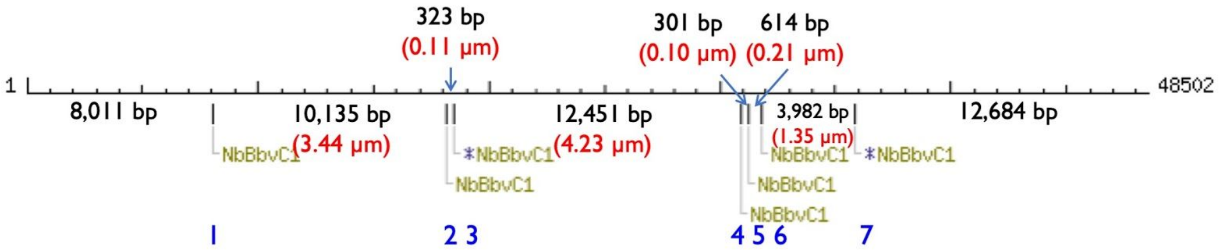


Figure S6: The location of nicking sites in $\lambda$DNA. The blue numbers reference the tag in the text. The black numbers show the distance in base pairs between two adjacent tags, with the red length indicating the calculated distance in μm.

Figure S6 shows the nicking sites location along the $\lambda$DNA. The NbBbvCI nicking enzyme locates the sequence of "CCTCA↑GC" and cut one strend. Then the biotinylated dNTP binds the nicking sites. So the mono-streptavidin tags are supposed to locate at the nicking sites. The distance between any two binding sites, from short to long, are $d_{45}$ is 301 bp : 102 nm, $d_{23}$ is 323 bp : 110 nm, $d_{56}$ is 614 bp : 209 nm, $d_{46}$ is 915 bp : 311 nm, $d_{67}$ is 3982 bp : 1354 nm, $d_{57}$ is 4596 bp : 1563 nm, $d_{47}$ is 4897 bp : 1665 nm, $d_{12}$ is 10135 bp : 3446 nm, $d_{13}$ is 10458 bp : 3556 nm, $d_{34}$ is 12451 bp : 4233 nm, $d_{35}$ is 12752 bp : 4336 nm, $d_{24}$ is 12774 bp : 4343 nm, $d_{25}$ is 13075 bp : 4446 nm, $d_{36}$ is 13366 bp : 4544 nm, $d_{26}$ is 13689 bp : 4654 nm, $d_{37}$ is 17348 bp: 5898 nm, $d_{27}$ is 17671 bp : 6008 nm, $d_{14}$ is 22909 bp : 7789 nm, $d_{15}$ is 23210 bp : 7891 nm, $d_{16}$ is 23824 bp : 8100 nm, $d_{17}$ is 27806 bp : 9454 nm.

We found the nicking sequence along the lambda DNA. So we learned the base pair distance between two adjacent sites. We used 0.34 nm/bp to calculate the distance between the tags. We show the distance in Figure S6.

#### 8. Examples of Scans

Figure S7 to Figure S14 show examples of scans. Cases where tags are visually present but went undetected by the code are marked as "missed."

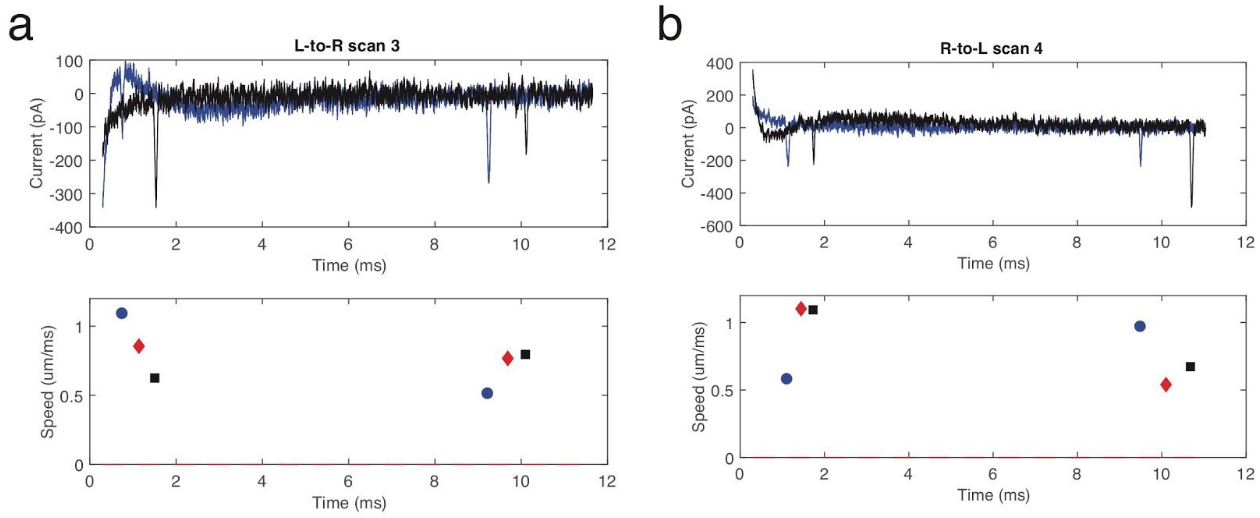


Figure S7: Representative examples of entry and exit signals from event (ii) in Table S2. Entry and Exit signals are blue and black, respectively, with detected tag speeds in the lower plot as a blue circle (Entry) or black square (Exit). The pore-to-pore speeds are shown as red diamonds (method of calculation described in the main text).

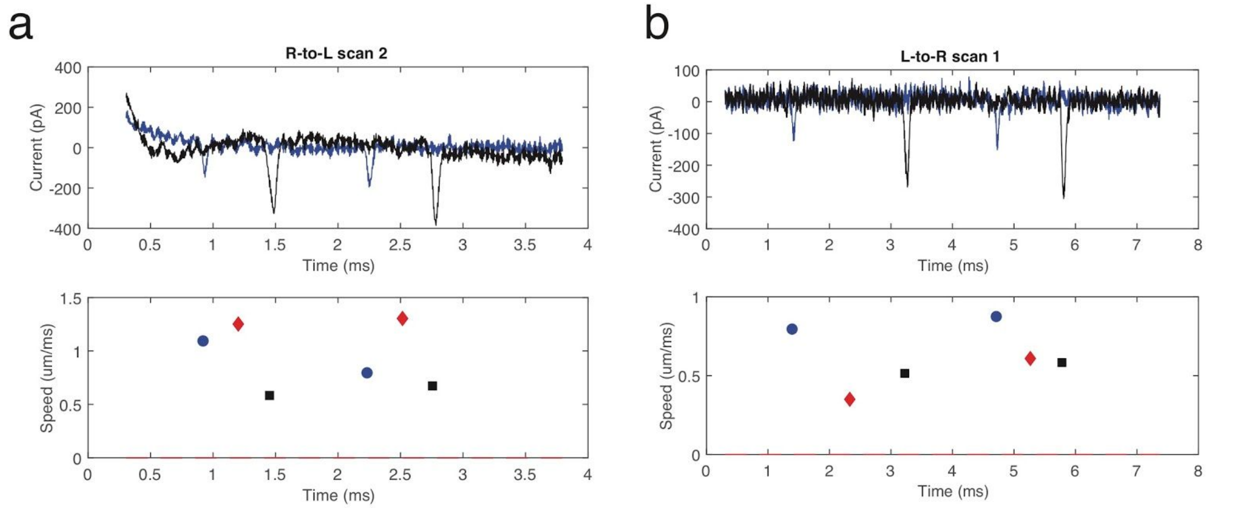


Figure S8: Representative examples of entry and exit signals from event (iii) in Table S2. Entry and Exit signals are blue and black, respectively, with detected tag speeds in the lower plot as a blue circle (Entry) or black square (Exit). The pore-to-pore speeds are shown as red diamonds (method of calculation described in the main text).

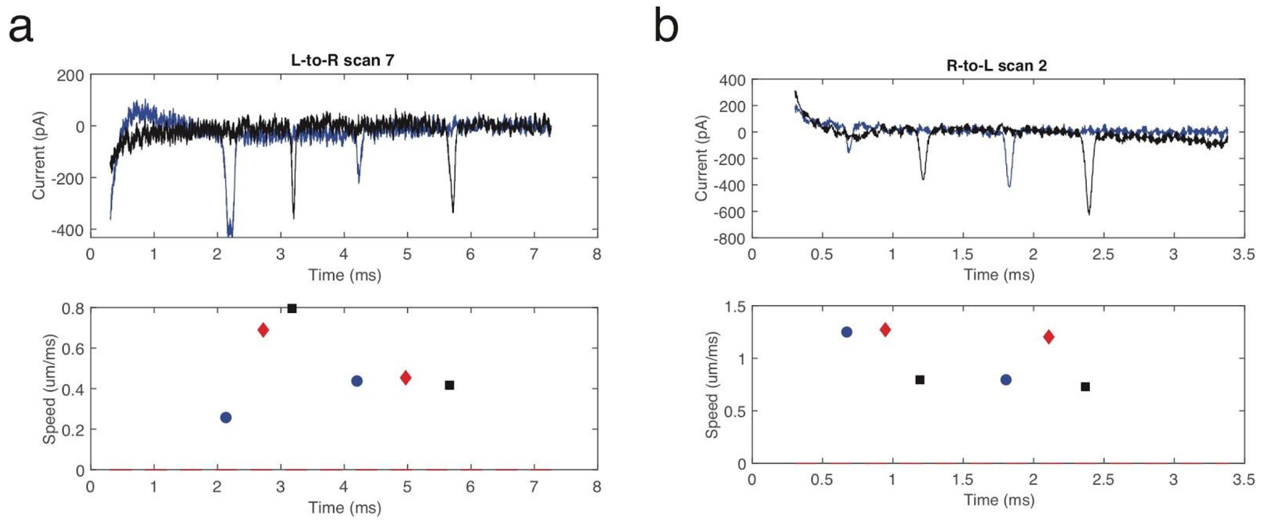


Figure S9: Representative examples of entry and exit signals from event (iv) in Table S2. Entry and Exit signals are blue and black, respectively, with detected tag speeds in the lower plot as a blue circle (Entry) or black square (Exit). The pore-to-pore speeds are shown as red diamonds (method of calculation described in the main text).

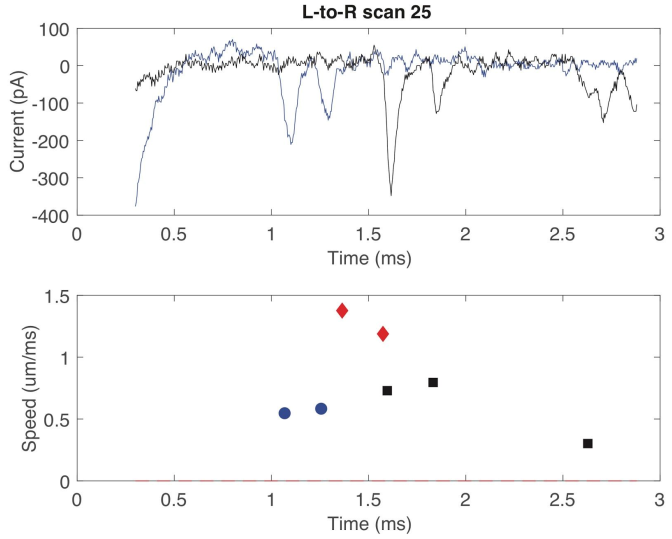


Figure S10: Representative examples of entry and exit signals from event (v) in Table S2. Entry and Exit signals are blue and black, respectively, with detected tag speeds in the lower plot as a blue circle (Entry) or black square (Exit). The pore-to-pore speeds are shown as red diamonds (method of calculation described in the main text).

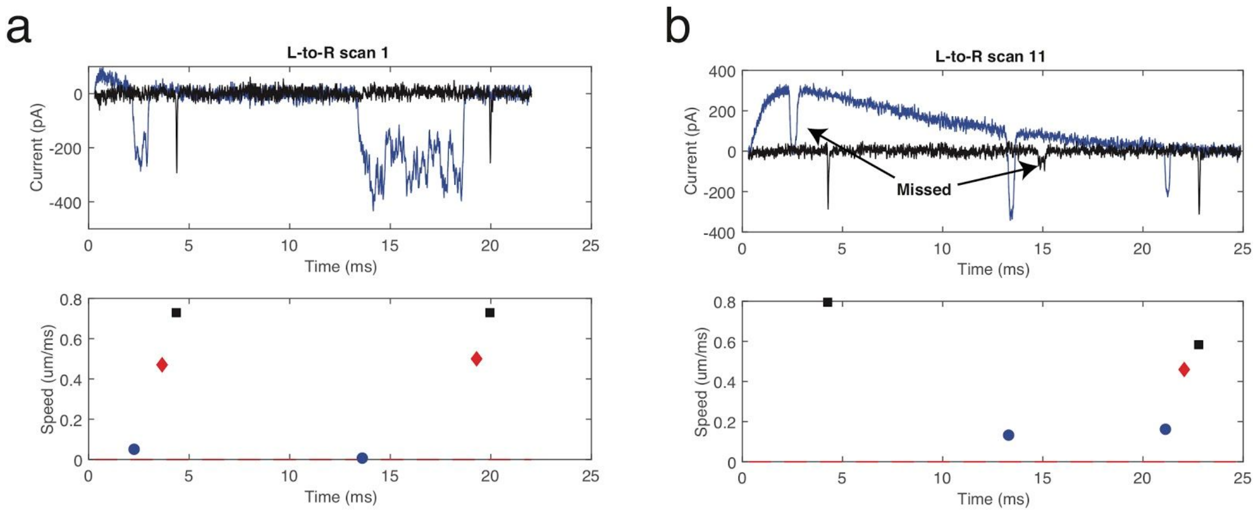


Figure S11: Representative examples of entry and exit signals from event (vi) in Table S2. Entry and Exit signals are blue and black, respectively, with detected tag speeds in the lower plot as a blue circle (Entry) or black square (Exit). The pore-to-pore speeds are shown as red diamonds (method of calculation described in the main text). Scan b shows a missed tag at the exit ($I_2$) resulting in a register-shift in the tag pair for subsequent scanning (also shown in main text Figure 4d-f).

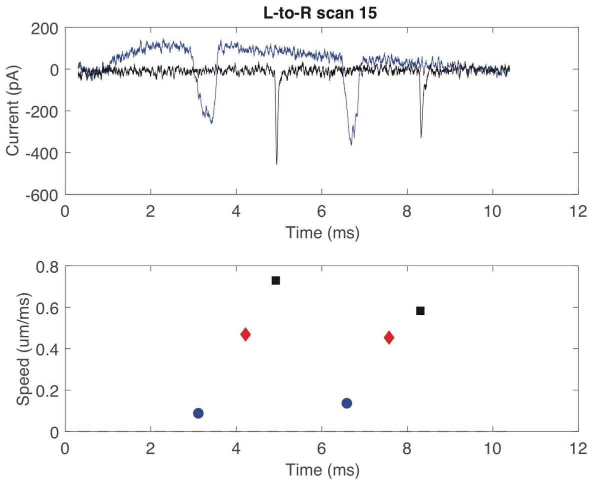


Figure S12: Representative examples of entry and exit signals from event (vii) in Table S2. Entry and Exit signals are blue and black, respectively, with detected tag speeds in the lower plot as a blue circle (Entry) or black square (Exit). The pore-to-pore speeds are shown as red diamonds (method of calculation described in the main text).

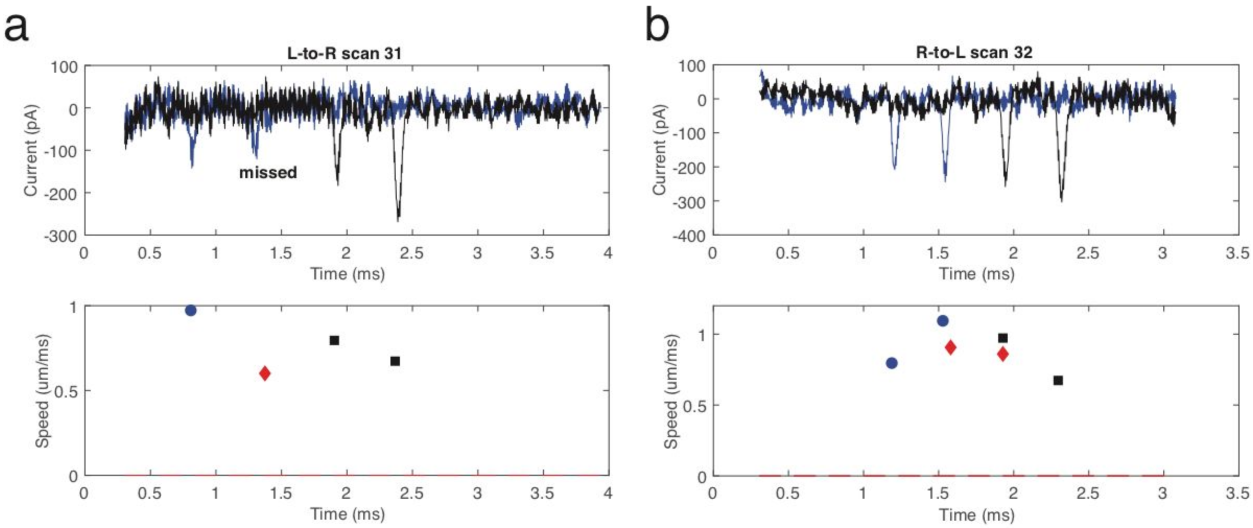


Figure S13: Representative examples of entry and exit signals from event (viii) in Table S2. Entry and Exit signals are blue and black, respectively, with detected tag speeds in the lower plot as a blue circle (Entry) or black square (Exit). The pore-to-pore speeds are shown as red diamonds (method of calculation described in the main text).

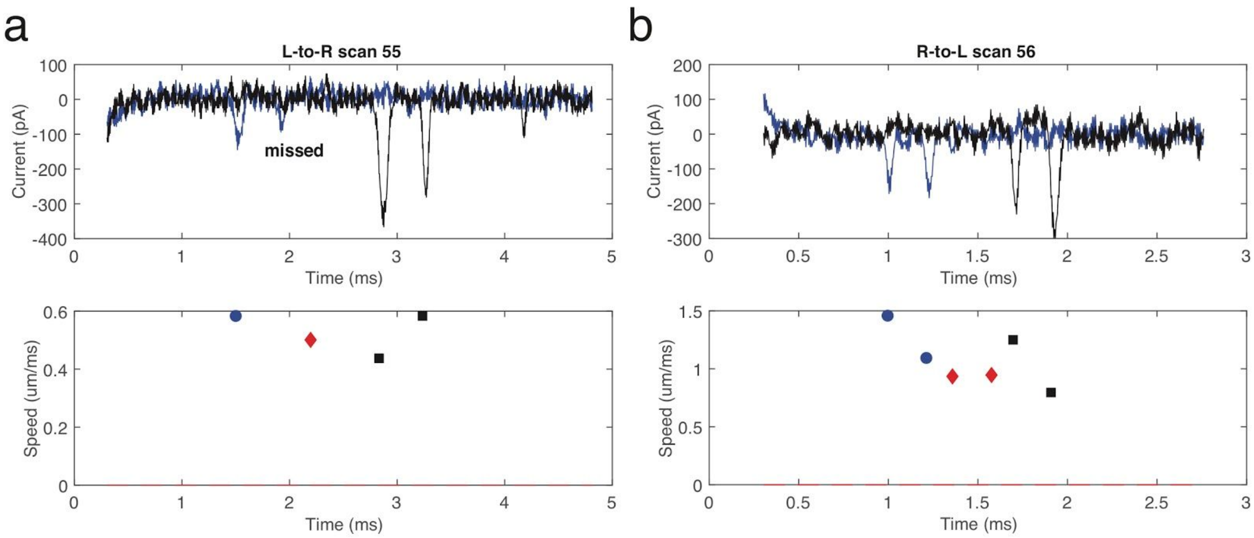


Figure S14: Representative examples of entry and exit signals from event (ix) in Table S2. Entry and Exit signals are blue and black, respectively, with detected tag speeds in the lower plot as a blue circle (Entry) or black square (Exit). The pore-to-pore speeds are shown as red diamonds (method of calculation described in the main text).

### 9. Tag Profile Characterization via Least-squares Fitting

In this section we describe our approach for obtaining peak-position and peak width at half-maximum using least-squares fitting of a model tag blockade profile. First, the raw multi-scan event is broken up into all component L-to-R and R-to-L scans. Then, the data is inverted (by reversing the sign of the current values) and Matlab's *findpeaks* algorithm is used to identify local maxima corresponding to the individual tag blockades (an example of the analysis protocol is shown in Figure S15). For pore 1 (entry pore for L-to-R polarity), voltages are changed upon termination of a scan to enable directional reversal, inducing capacitance transients in the current. These transients xacare fit to an exponential model (Figure S15a). Note that, prior to fitting the transient, a fixed region of $400\,\mu s$ around each identified tag is removed to ensure the tag blockades do not interfere with the background fit. The fitted transient background for the pore 1 channel is removed and then each tag blockade is fitted to a model profile (Figure S15b). While the transient is mostly removed, there is a small residual within about 1 ms of the scan start, as the exponential is not an exact model. There is no risk of mistaking this residual as a tag blockade, as it corresponds to a current increase above baseline and occurs in the same location for each rescan.

This model profile has the following functional form:

$$I(t) = I_{b1} + I_{b2}t - \frac{I_o}{2}\left[erf\left(\frac{t-(t_o-\Delta t/2)}{\sqrt{2}\sigma}\right) - erf\left(\frac{t-(t_o+\Delta t/2)}{\sqrt{2}\sigma}\right)\right] \quad \text{S2}$$

This form is based on the convolution of a box of width $\Delta t$ and height $I_o$ with a normalized Gaussian of width $\sigma$. In limit that $\Delta t >> \sigma$ this model has the form of a broadened box; in the limit $\Delta t << \sigma$ it has the form of a Gaussian function. For the purposes of tag-profile characterization, this model has the advantage that it can describe both tag transits of long duration with a broadened box shape and rapid tag transits with a more peaked shape. The width at half maximum can be obtained as a function of $\Delta t$ and $\sigma$ (in the limit $\Delta t >> \sigma$, the width is $\Delta t$, in the limit $\sigma << \Delta t$ the width is that of a Gaussian function, in the intermediate case, the width at half-maximum is computed numerically as a function of $\Delta t / \sigma$ and then interpolation used to obtain the width from any combination of $\Delta t$ and $\sigma$). In addition, we have found that the fitting is more robust if we add a linear function to account for any residual background variation that is not captured by the exponential fit. In practice, this fitting is performed using Matlab's *lsqcurvefit* for each tag over a range of fixed duration (typically ~ 0.5-1 ms) centered on the location of the preliminary tag position identified by *findpeaks*. The parameters determining the linear background ($I_{b1}$ and $I_{b2}$) are determined by averaging the background over $60\,\mu s$ at the beginning and end of the tag-centered interval (in cases where the background is flat, note that $I_{b2} \approx 0$). For pore 2 (exit pore on L-to-R) the same procedure is applied, except there is no need to remove a capacitance transient as the voltage at pore 2 is held constant (Figure S15c). Note that the model can accommodate both the peaked Gaussian shape of the tag blockades in Figure S15c and the flatter box-like shape of the blockades in Figure S15b.

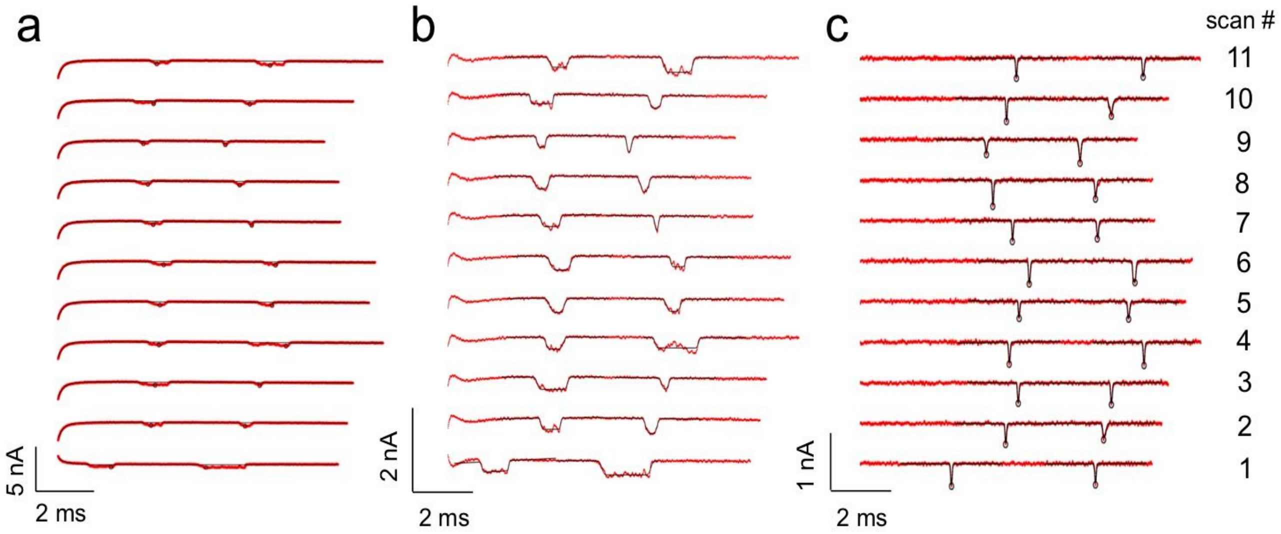


Figure S15: Peak Fitting Approach for Tag Profile Characterization. Here we illustrate our least-squares fitting approach for finding the center position and width at half-maximum of tag blockade profiles. (a) Example of multi-scans from pore 1 (entry) with L-to-R polarity (red). These scans are fitted with an exponential model (black bold curve) to remove the capacitance transient. The open circles show initial detected peak-positions using a local peak-finding algorithm. (b) Scans from (a) with capacitance transient removed. The bold curves at each peak are fits obtained with our erf-function model function for the tag blockade profile. (c) Example of multi-scans from pore 2 (exit) with L-to-R polarity (red). The open circles show initial detected peak-positions using a local peak-finding algorithm; the bold-curves at each peak are fits obtained with the erf-model. This event was taken with a device that had a 0.66 μm pore-to-pore spacing, pore 1 had a diameter of 23 nm and pore 2 had a diameter of 23 nm (Table S1). A voltage of 150 mV was applied to pore 1 and a voltage of 300 mV was applied to pore 2 during the L-to-R scans.

#### 10. Tag Alignment Procedure

Here we describe the details of the tag alignment algorithm. Each successive scan represents a measurement of an underlying binding pattern of tags over a certain region of the molecule. The scans in each series of fixed polarity (i.e. L-to-R or R-to-L) have a relative translational offset, arising from the fact that different portions of the molecule are observed in each scan. There is also stochastic variation in tag positions, arising from Brownian fluctuations (see Figure S18a,e, these events correspond to results shown in manuscript Fig. 5). These effects complicate correct association of tags across multiple scans (i.e. how do we ensure that a tag observed in scan i corresponds to the same tag in scan j, with “same” implying that the tags correspond to a single tag at the same sequence position?). The objective of this algorithm is to introduce a systematic procedure for aligning scans in a given series (L-to-R or R-to-L for pore 1 or pore 2) by identifying the correct corresponding tag pairs between successive scans and then removing translational offset between the scan pairs.

The core of the algorithm is a function, *pairalign*, that computes a measure of alignment error based on the squared difference between the $i^{th}$ and $j^{th}$ scans in a given scan set. This function assumes that at least two tags are shared, or are `common,' between the $i^{th}$ and $j^{th}$ scan. In order to show that this assumption is valid for our multi-scan data, the spacing between the tags in each scan can be computed and compared. What we observe typically are cases where the spacing in each scan fluctuates around a fixed mean value (see Figure S16b), or occasionally bimodal situations where two mean spacing values are observed (see Figure S16f). In bimodal situations, the spacings often exchange at a scan where three trags are observed. Thus, it is reasonable to assume that two common tags will be present between successive scan pairs (scans i and j=i-1).

In order to identify the common tags in a scan pair, *pairalign* computes a measurement of alignment error over all potential alignments of the two scans. Each of these potential, or test alignments, is determined by choosing a tag pair between scan i and j and then shifting scan i by the correct time interval to bring this chosen tag pair into alignment (we will call this tag pair the "aligned pair"). Then, omitting the aligned pair, the squared distance is computed for all possible tag pairs that can be formed between the scans. The list of these possible pairings is sorted by squared difference and the distinct pairings with minimum squared difference are obtained (note that the number of distinct pairs is equal to $\min(n_{tag,i}, n_{tag,j})$ where $n_{tag,i}$ is the number of tags on the $i^{th}$ scan and $n_{tag,j}$ is the number of tags on the $j^{th}$ scan). The overall alignment error, for a given choice of aligned pair, is the sum of the squared differences of these distinct pairings with minimum squared difference. The aligned pair taken as the correct common tag between the two scans is the aligned pair that yields the minimum overall alignment error. As a consequence of this procedure, which identifies a set of pairings between tags in the scan, a correspondence table can be constructed between the tags in scan i and scan j, yielding all common tag pairs shared between the scans.

Blue circles are spacings for scans with only two tags observed (for which there is only one spacing). Red squares and magenta triangles represent the two distinct nearest neighbor spacings when three tags are observed. (g) Aligned tag positions; meaning of data color and shape scheme same as in (e). (h) Aligned and distance calibrated tag positions: the data color and shape scheme now reflects the group assignments and corresponds to true physical tags (tag A, blue circles; tag B, red squares and tag C magenta triangles). Black circles are final averaged tag positions corresponding to tags A, B and C. Note that the first scan corresponds to an alignment with an error over threshold and is removed from computation of the averaged tag position (indicated by cross).

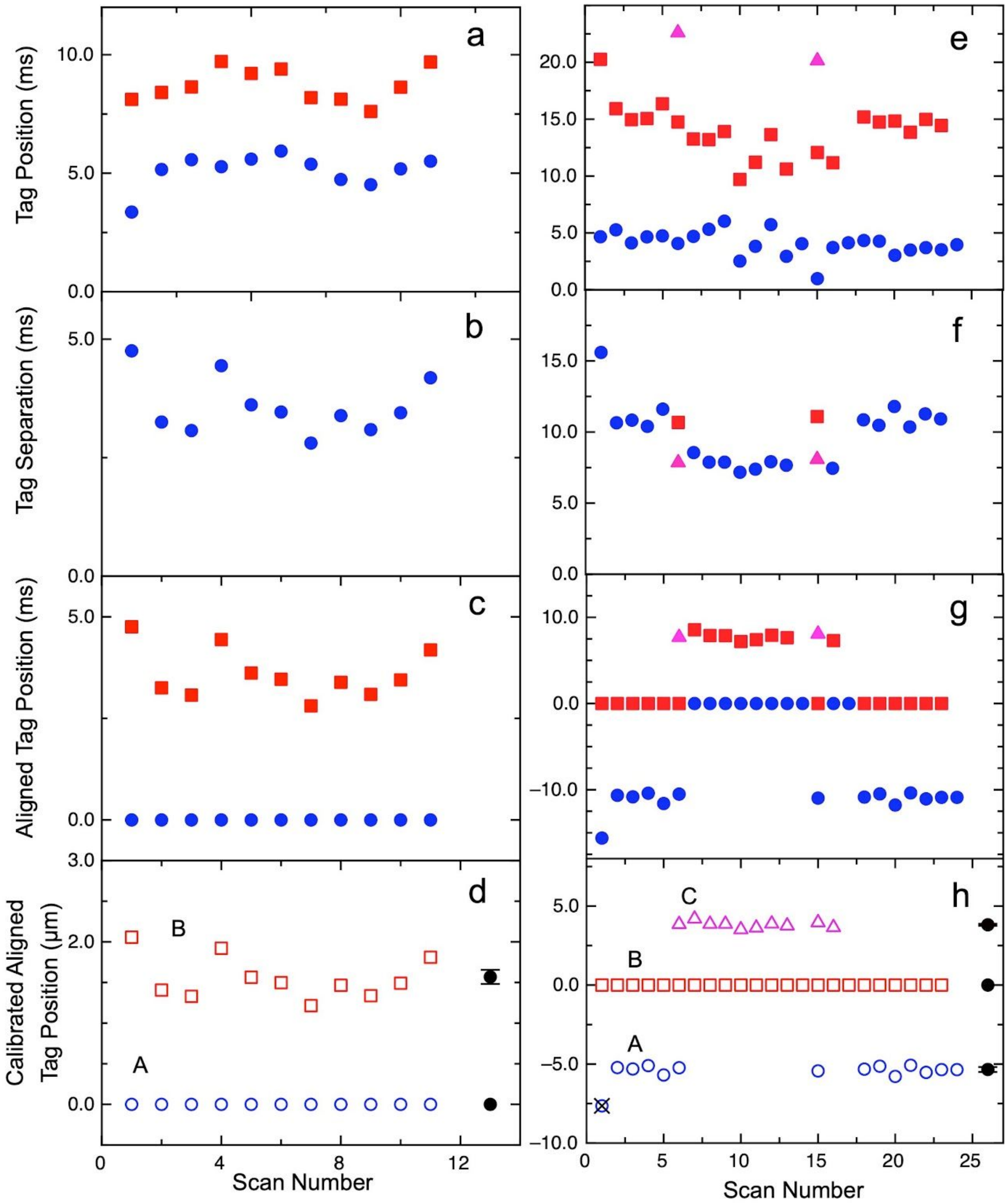


Figure S16: Tag alignment procedure. (a) Example of tag-position versus scan number for an event with two tags (blue circles, tag measured closest to scan start in each scan, red squares tag observed furthest from scan start). (b) Spacing between tags as a function of scan number for event in (a). (c) Aligned tag positions for event in (a). The distance calibrated and aligned tag position for event in (a) with final averaged tag positions (here blue circles correspond to associated measurements for tag A, red squares correspond to associated measurements for tag B). (e) Example of tag-position versus scan number for an event with three tags (blue circles, tag position closest to event start, red squares tag at intermediate distance, magenta triangles tag observed furthest away from event start). (f) Spacing between tags for event in (e).

*Pairalign* is applied between successive scan pairs in a scan set. The correspondence table, applied iteratively, enables association of tags observed in each scan into groups that correspond to the true physical tags present on the molecule (Figure S16d,h). The initial number of groups corresponds to the number of tags in the first scan. If scan i has more tags than scan j, then new groups are introduced.

A problem occurs when a scan has only one tag. If scan i has one tag, then the common tag in j=i-1 is chosen as the tag with minimum separation to the one tag in scan i. If scan j=i-1 has only one tag, then the algorithm seeks to bring the $i^{th}$ scan into alignment with the i-2 scan. Another problem is that tags corresponding to different groups can be inadvertently associated together, particularly if a third tag is missed in the scan bordering the groups. The code seeks to prevent this in the following way. If the overall alignment error of the $i^{th}$ and i-1 scans is greater than a preset threshold, yet the alignment to successive tags is below threshold, then this indicates a junction between two different tag groups that are incorrectly grouped together. The code will check the alignment between the $i^{th}$ scan and scans preceding the i-1 scan that yielded the large alignment error. All alignments that also yield an error over the threshold value are assumed to belong to a different group and assigned a new group number. Groups which correspond to only one scan are further checked by associating the scan with all other scans; if alignments are found yielding an error below the threshold value than these groups are consolidated (example is scan 15 in Figure S16h). If a single over-threshold scan cannot be associated with additional scans, i.e. is a group corresponding to only one scan, it is removed from the analysis as an outlier (an example is the first scan for event shown in Figure S16h).

At this point the largest group is defined to be the origin tag. All scans are shifted by the amount required to bring the origin tag to zero. For a scan i that does not contain the origin tag, the scan is associated through a common tag pair to a scan j that does contain the origin tag. The scan i is then shifted to bring the common tag to the same position as in scan j. This step removes residual translational offset.

Lastly, a final single molecule barcode is constructed by averaging together all tag measurements that belong to a given group (Figure S16d,h). The error on a given tag location can be obtained as the standard-deviation of the mean for the group.

#### 11. Tag-to-Tag Separation Statistics from Nine Multi-Scan Events

Table S3 shows data on nine different multi-scan events, five of which are summarized in the main text as Table 1 (unique cycle numbers reveal the correspondence). Observe that more data can come from one pore versus the other, or can be balanced in volume across both pores. One of the two scan directions will also produce more data than the other, and this appears to be device dependent and/or voltage-setting dependent. By example for pore 2 data, (iii) and (iv) are from one experiment that produced more data L-to-R than R-to-L, while (viii) and (ix) are from another experiment that produced more data R-to-L than L-to-R.

Table S3: **Statistics related to tag separation estimation from nine multi-scan events**

| Event & No. Cycles$^\sharp$ | Scan Dir.$^\ddagger$ | Entry Tag Speed ($\mu$m/ms)$^\dagger$ | Exit Tag Speed ($\mu$m/ms)$^\dagger$ | Pore-to-Pore Speed ($\mu$m/ms)$^\dagger$ | Pore 2 Tag-to-Tag Separations ($\mu$m)$^*$ | Combined Tag-to-Tag Separations ($\mu$m)$^*$ |
|---|---|---|---|---|---|---|
| (i), 48 | L-to-R | $0.86 \pm 0.18$ (72) | $0.65 \pm 0.21$ (101) | $0.35 \pm 0.29$ (41) | a. $0.074 \pm 0.0068$ (23) | a. $0.073 \pm 0.0051$ (39) |
| | | | | | b. $0.77 \pm 0.046$ (2) | b. $1 \pm 0.078$ (17) |
| | R-to-L | $0.92 \pm 0.25$ (94) | $0.75 \pm 0.33$ (88) | $0.81 \pm 0.16$ (57) | a. $0.1 \pm 0.0049$ (32) | a. $0.12 \pm 0.0063$ (56) |
| | | | | | b. — (0) | b. $1.4 \pm 0.031$ (21) |
| (ii), 27 | L-to-R | $0.63 \pm 0.19$ (27) | $0.82 \pm 0.21$ (48) | $0.73 \pm 0.22$ (26) | $6.6 \pm 0.28$ (21) | $6.6 \pm 0.25$ (24) |
| | R-to-L | $0.77 \pm 0.26$ (41) | $0.81 \pm 0.26$ (27) | $0.81 \pm 0.15$ (27) | $9.3 \pm 0.77$ (20) | $8.9 \pm 0.62$ (26) |
| (iii), 30 | L-to-R | $0.64 \pm 0.19$ (50) | $0.43 \pm 0.21$ (60) | $0.48 \pm 0.25$ (49) | $1.5 \pm 0.12$ (28) | $1.5 \pm 0.12$ (48) |
| | R-to-L | $1.2 \pm 0.22$ (39) | $0.82 \pm 0.19$ (47) | $1.2 \pm 0.15$ (38) | $1.5 \pm 0.026$ (15) | $1.5 \pm 0.017$ (38) |
| (iv), 70 | L-to-R | $0.53 \pm 0.21$ (118) | $0.52 \pm 0.21$ (138) | $0.54 \pm 0.17$ (118) | $1.3 \pm 0.045$ (68) | $1.3 \pm 0.034$ (116) |
| | R-to-L | $0.94 \pm 0.3$ (102) | $0.76 \pm 0.22$ (134) | $1.1 \pm 0.17$ (99) | $1.4 \pm 0.017$ (32) | $1.3 \pm 0.011$ (96) |
| (v), 38 | L-to-R | $0.51 \pm 0.24$ (74) | $0.66 \pm 0.24$ (92) | $1.2 \pm 0.2$ (73) | a. $0.23 \pm 0.0087$ (38) | a. $0.24 \pm 0.0057$ (72) |
| | | | | | b. $1.0 \pm 0.078$ (13) | b. $1.0 \pm 0.078$ (13) |
| (vi), 24 | L-to-R | $0.19 \pm 0.14$ (39) | $0.6 \pm 0.19$ (47) | $0.46 \pm 0.14$ (35) | a. $4 \pm 0.1$ (9) | a. $4 \pm 0.073$ (17) |
| | | | | | b. $5.2 \pm 0.038$ (9) | b. $5.2 \pm 0.038$ (9) |
| (vii), 11 | L-to-R | $0.13 \pm 0.094$ (23) | $0.62 \pm 0.15$ (22) | $0.41 \pm 0.11$ (21) | $1.4 \pm 0.036$ (8) | $1.4 \pm 0.024$ (16) |
| (viii), 31 | L-to-R | $0.91 \pm 0.3$ (12) | $0.75 \pm 0.12$ (22) | $0.93 \pm 0.41$ (12) | $0.31 \pm 0.022$ (9) | $0.31 \pm 0.022$ (9) |
| | R-to-L | $0.94 \pm 0.16$ (58) | $0.76 \pm 0.12$ (57) | $0.99 \pm 0.19$ (56) | $0.3 \pm 0.005$ (28) | $0.31 \pm 0.0039$ (55) |
| (ix), 65 | L-to-R | $0.68 \pm 0.19$ (38) | $0.66 \pm 0.2$ (74) | $0.71 \pm 0.25$ (38) | $0.2 \pm 0.0083$ (18) | $0.2 \pm 0.009$ (20) |
| | R-to-L | $0.96 \pm 0.17$ (122) | $0.79 \pm 0.15$ (128) | $0.95 \pm 0.16$ (121) | $0.21 \pm 0.0025$ (57) | $0.21 \pm 0.0021$ (119) |

$^\sharp$ Events (i-iv), (v), (vi-vii) and (viii)-(ix) are from common chips E, G, A and F (Table S1), respectively.
$^\ddagger$ R-to-L not reported where an insufficient number of tags were detected.
$^\dagger$ Mean $\pm$ standard deviation (count). No data was trimmed.
$^*$ Mean $\pm$ standard deviation of the mean (count). One or two (a,b) separations are reported where two or three tags, respectively, were reliably detected.

The pore-to-pore speed coefficient of variation (CV, equal to standard deviation divided by the mean) provides a metric that can be used to assess how reliable the distances estimates are, since its a measure of heterogeneity of motion from scan to scan. The pore-to-pore speed CV is high at 83% for the L-to-R scans and more reasonable at 20% for the R-to-L scans for event (i) (also event (i) in main text Table 1). As described in main text Figure 3, only the shorter tag-pair distance estimates were available in $I_2$ for event (i), and so the second longer tag-pair spacing (B-C in Figure 3) is averaged and reported in Table S3 using only data from $I_1$.

We can match estimates to mappable tag-to-tag distances using the map in Figure S7. For event (i), ignoring the high speed CV data that is L-to-R, the R-to-L estimates that have 120 nm and 1400 nm adjacent are tags 4-5 (100 nm) and 5-7 (1560 nm). For event (ii), again using the scan direction R-to-L with lower speed CV, the estimate of 8900 nm is closest to tags 1-7 which is the largest inter-tag gap at 9450 nm. These two events, (i) and (ii), therefore cover the smallest and largest inter-tag gaps for our model reagent, from 300 bp to 27,800 bp. Event (iii) and (iv) are consistent with 6-7 (1350 nm) and 5-7 (1563 nm). Event (v) is consistent with adjacent tag pairs of 5-6 (210 nm) and 6-7 (1350 nm), though the estimate underpredicts the latter badly. Event (vi) is consistent but apparently systematically overpredicts the adjacent tag pairs of 1-2 (3440 nm) and 2-5 (4450 nm), while event (vii) is closest to 6-7 (1350 nm). Lastly, events (viii) and (ix) are closest to 4-6 (310 nm) and 5-6 (210 nm), respectively.
